# Transcriptional control of *C. elegans* male tail tip morphogenesis by DMD-3

**DOI:** 10.1101/2024.12.19.629486

**Authors:** Porfirio Fernandez, Sevinç Ercan, Karin C. Kiontke, David H. A. Fitch

## Abstract

Sexual dimorphic morphogenesis is governed by DM-domain transcription factors (TFs) in many animals, but how these transcriptional control links to the morphogenetic mechanisms is insufficiently known. The DM-domain TF DMD-3 in *C. elegans* is the master regulator of a male-specific development that changes the shape of the tail tip from long and pointed in larvae to short and round in adults. This tail tip morphogenesis (TTM) requires cell-shape changes, cell migration and fusion. To understand how transcriptional regulation by DMD-3 governs TTM, we used male-specific ChIP-seq to identify its direct targets. We found 1,755 DMD-3 bound sites. We identify a DMD-3 associated binding motif and validate its function in TTM. This motif is similar to the binding motif of EOR-1, and we suggest that DMD-3 acts cooperatively with EOR-1 and possibly other TFs. DMD-3 targets 270 genes that play a role in TTM. These genes include other TFs but also effectors and components of morphogenetic mechanisms. By deleting the DMD-3 bound region endogenously and observing changes in reporter expression and tail tip phenotypes, we identify tissue specific enhancers in the cis-regulatory region of *fos-1, pan-1, nmy-2* and *hmr-1* that play a role in TTM. For *fos-1*, we propose that a feed-forward loop is responsible for tail-tip specific increase of gene-expression. This study provides insights into the architecture of the genetic regulatory network controlling a morphogenetic process.

**Article Summary:** DM domain transcription factors are often responsible for sexually dimorphic morphogenesis, but how they connect to morphogenetic mechanisms is insufficiently known. Here, we use ChIP-seq to determine the direct targets of DMD-3, which is the master regulator of male-specific tail tip morphogenesis (TTM) in *C. elegans*. We find that DMD-3 targets 270 TTM genes which include other transcription factors but also effectors and components of morphogenetic mechanisms. This study provides insights into the architecture of the genetic regulatory network controlling a morphogenetic process.

## Introduction

Morphogenesis, the creation of biological form, requires precise spatial and temporal coordination of cell behaviors such as migration, shape change or fusion. How the processes during morphogenesis are regulated by gene-regulatory networks is insufficiently known (e.g. Gill et al.). Most studies focus on embryogenesis, however, morphogenesis also happens post-embryonically, e.g. to create differences in morphology between juveniles and adults and between males and females. Across animals, from corals to mammals, DM-domain-related transcription factors (DMRTs) are involved in sexual development (Matson and Zarkower 2012; Bellefroid et al. 2013; Traylor-Knowles et al. 2015). In many systems, they drive male-specific cell fates (Kopp 2012; Smith et al. 2024). One DMRT that regulates male-specific development in *C. elegans* is DMD-3. DMD-3 is involved in the development of the somatic gonad (Smith et al. 2024), male-specific neurons (Serrano-Saiz et al. 2017), the copulatory structures in the tail and the tail tip (Mason et al. 2008). In Hermaphrodites, *dmd-3* expression is repressed in all tissues except for the anchor cell by TRA-1, a TF downstream in the sex determination pathway (Mason et al. 2008).

Here, we study DMD-3 in the context of the tail tip, which is long and pointed in larvae and hermaphrodites but short and round in adult males. This difference arises during the last larval stage, when the four cells of the tail tip change shape, round up, migrate, and fuse (Fig. 1). This tail tip morphogenesis (TTM) is a simple and accessible model for morphogenesis and has been studied at the morphological, molecular, and transcriptional level (Nguyen et al. 1999; Mason et al. 2008; Nelson et al. 2011; Kiontke, Fernandez, et al. 2024; Kiontke, Herrera, et al. 2024).

**Figure 1.**
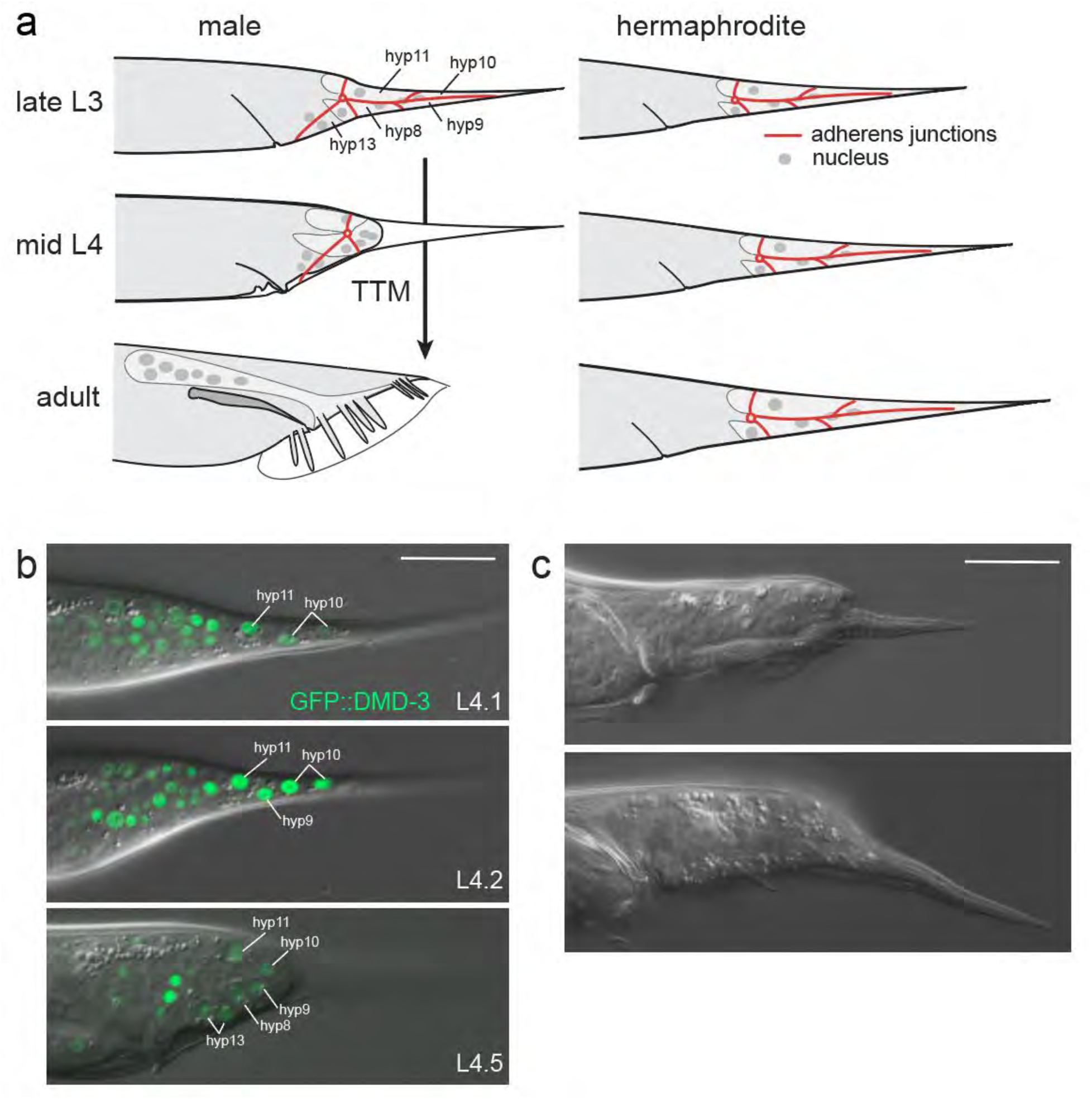
Wild-type tail tip morphogenesis (TTM) and DMD-3 expression. a) The *C. elegans* tail tip consists of 4 cells (hyp8-11) and is pointed in larvae of both sexes (early L4). In males during L4, adherens junctions (red) disassemble, the tail tip tissue dissociates from the cuticle, the tip rounds up, and the cells migrate anteriorly. The adult tail tip is short and round; the tail tip tissue has migrated anterior of the cloaca. Hermaphrodite tail tips remain pointed throughout development. b) Expression of DMD-3 endogenously tagged with GFP at the beginning of L4 when it has just begun in all tail tip cells (top), at the collection time for ChIP-seq when it is at its peak (middle), and when it is turning off after the rounding of the tail tip is completed (bottom). c) Examples of the adult tail phenotype of *dmd-3(0)* mutant males; TTM failed and the tail tip remains pointed. Scale bars = 20 µm.

TTM fails In *dmd-3(-)* mutants (Fig. 1), and misexpression of *dmd-3* in the tails of hermaphrodites is sufficient to cause ectopic TTM (Mason et al. 2008). DMD-3 is thus the master regulator for TTM. Genes that are involved in TTM upstream and downstream of DMD-3 were identified by mutagenesis screens and a whole genome RNAi screen (Zhao et al. 2002; Del Rio-Albrechtsen et al. 2006; Nelson et al. 2011; Herrera et al. 2016; Kiontke et al. 2019). From this work, it was proposed that the gene regulatory network for TTM has a bow-tie-shaped architecture with DMD-3 at the core (Nelson et al. 2011). Inputs that determine when during development, in which body part, and in which sex TTM is to occur feed into the core, which then regulates the mechanisms executing the processes of morphogenesis. A tissue-specific RNA-seq study comparing tail tips of wild-type males to those of hermaphrodites and *dmd-3(-)* males identified genes that are regulated by DMD-3 (Kiontke, Herrera, et al. 2024), adding more known components to the output part of the network downstream of the core, including a large number of other transcription factors. However, how exactly DMD-3 functions as the master regulator of TTM, how it is connected to its downstream network, and how this network is organized is still largely unknown.

In an extreme view, the function of a master regulator of morphogenesis could be to exclusively target other TFs which then regulate all genes required for the morphogenetic process. In such a network structure, the master regulator sits at the top of a regulatory hierarchy that controls most or all of the regulatory activities of the other transcription factors and the associated genes (Sikdar and Datta 2017). Alternatively, the master regulator could also target other genes. Such genes could encode effectors (e.g. small GTPases or protein modification enzymes) or key components of the cellular machinery for morphogenesis (e.g. cytoskeleton, vesicular transport, polarity). Examples of effector targets of a master regulator are the small GTPase RhoD/F as target of FoxF in Ciona trunk ventral cell migration (Christian …) and Fog, the ligand in a signaling pathway required for gastrulation in *Drosophila,* as a direct target of Twist (reference). Twist is also the master regulator for epidermal to mesenchymal transition in cancer metastasis, where it directly targets and downregulates E-cadherin (Vesuna et al. 2008).

To find the direct targets of the master regulator of TTM, we performed a male-specific whole-worm DMD-3 ChIP-seq experiment (chromatin immunoprecipitation followed by next-generation sequencing). This experiment found 1,755 DMD-3-bound sites near 6061 candidate target genes. We also identified a sequence motif significantly enriched at most of these sites, which we call the DMD-3-associated motif. We compared the candidate target genes obtained from whole worms with genes that were differentially expressed in the tail tip-specific RNA-seq study (Kiontke, Herrera, et al. 2024). We thus found 270 genes that act in the tail tip and are directly regulated by DMD-3.

The direct targets of DMD-3 are enriched for transcription factors but also contain members of many other functional classes, e.g. signaling molecules and components of the cytoskeleton, cell-cell junctions and the extracellular matrix. This suggests that the master regulator of TTM lies indeed at the top of a TF hierarchy, but it also directly regulates many other types of genes. We validated three of these genes as bona fide targets by deleting the DMD-3 binding sites in their 5’-regulatory regions. This analysis also revealed the existence of several tissue-specific enhancer elements in the cis-regulatory region of four genes.

## Materials and methods

### Strains

Strains were cultured on NGM plates seeded with *E. coli* OP50-1 under standard conditions (Stiernagle 2006). All strains investigated here contained the *him-5(e1490)* mutation or an identical allele generated by CRISPR genome editing.

**Table 1.**
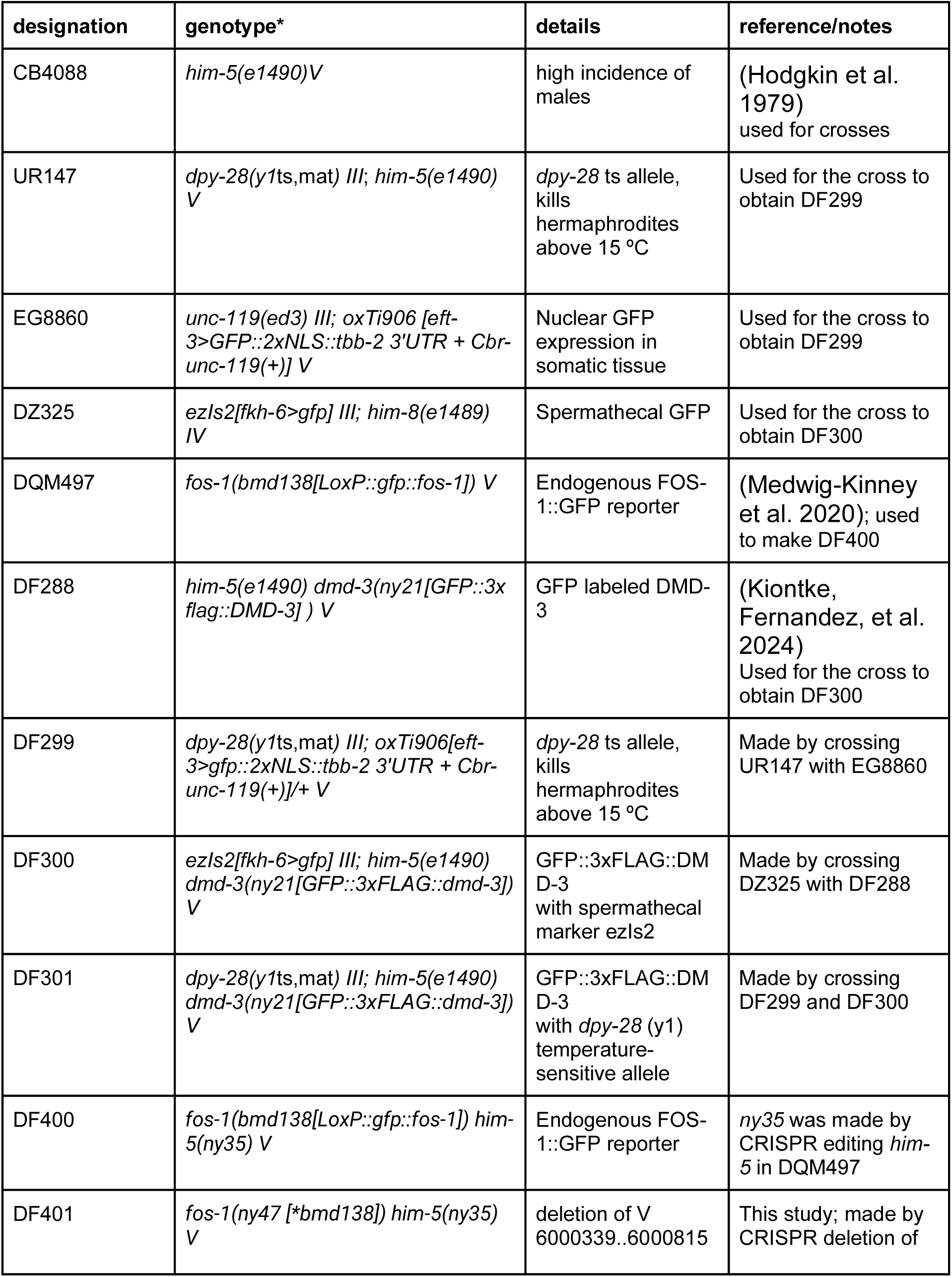
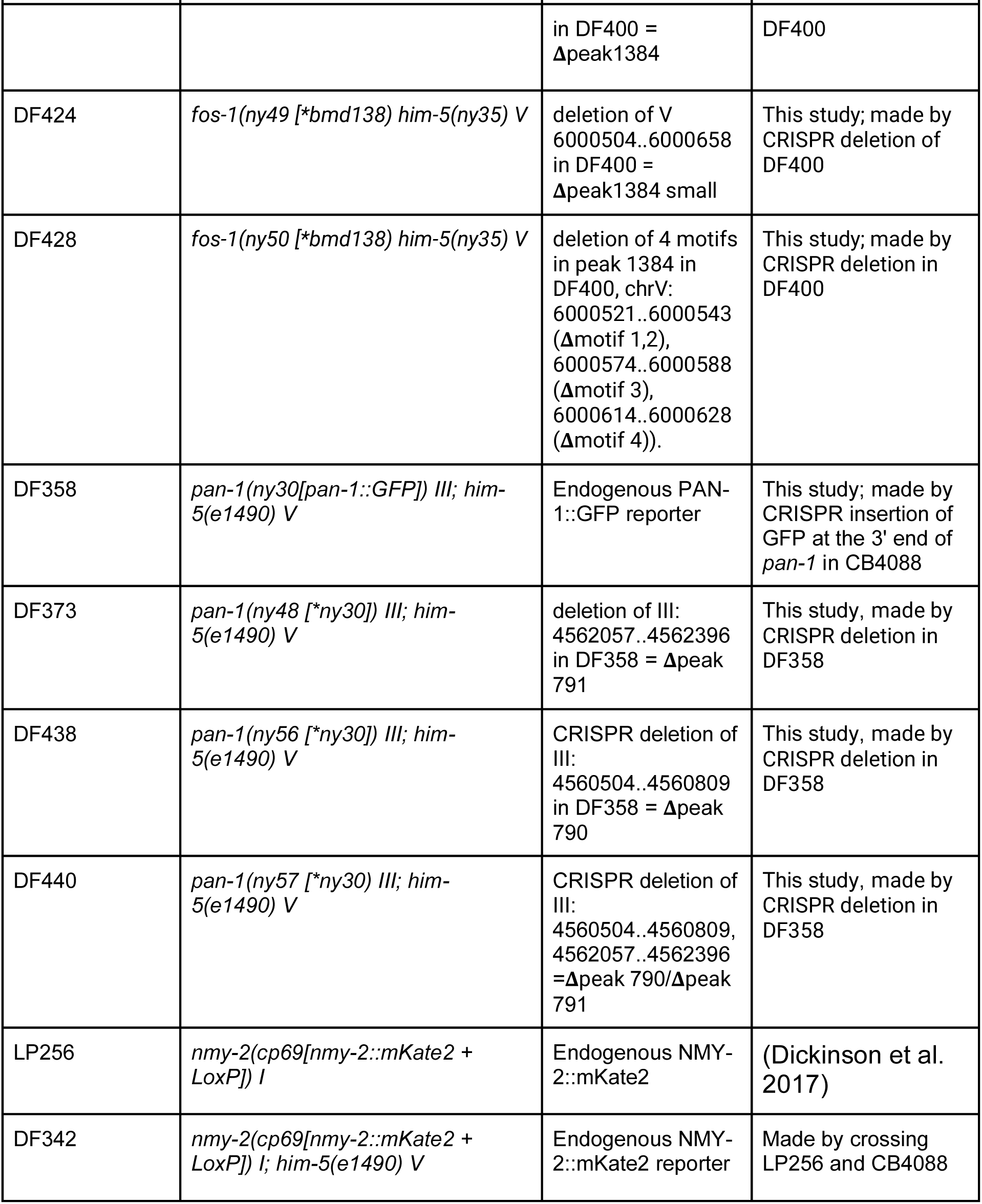
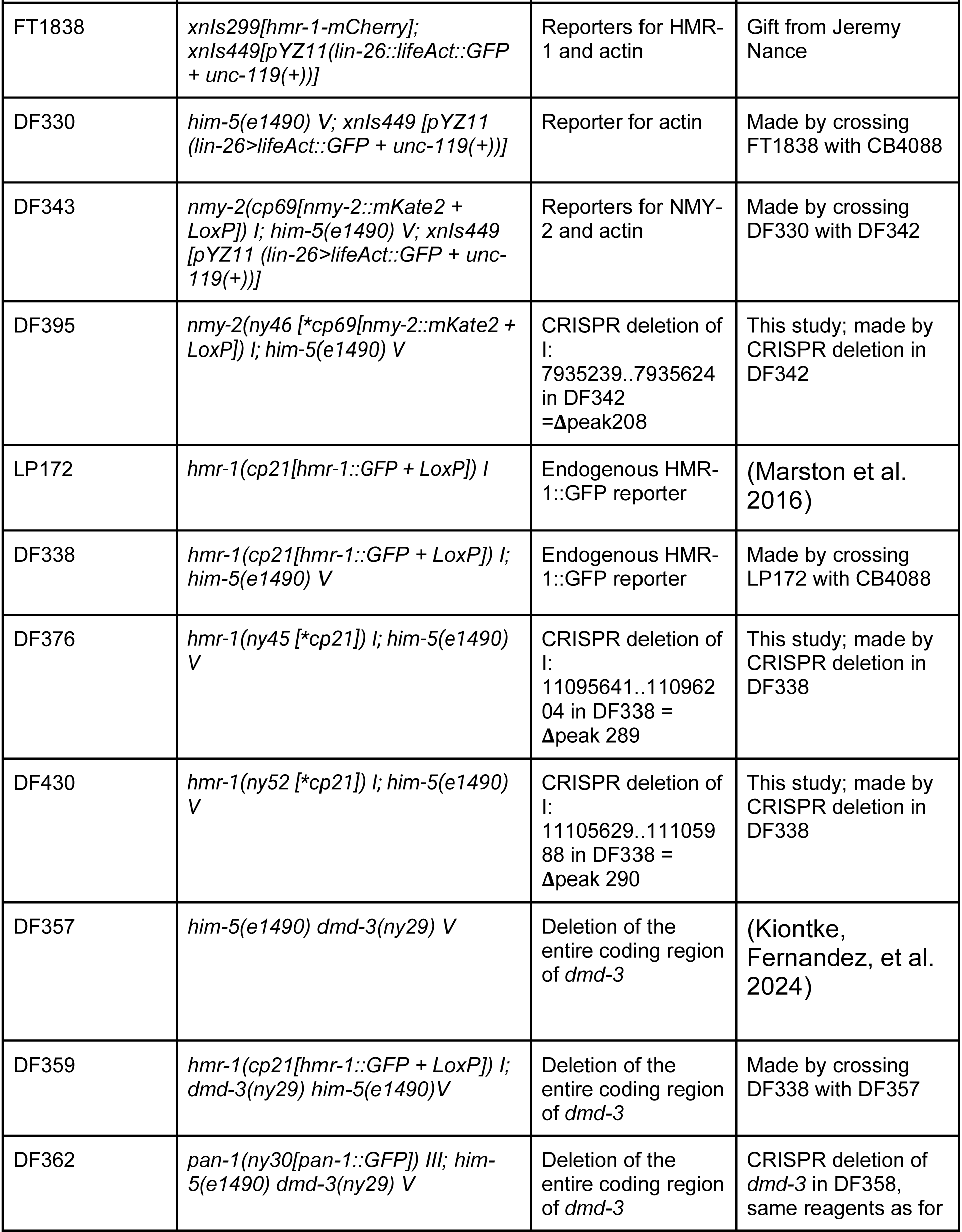
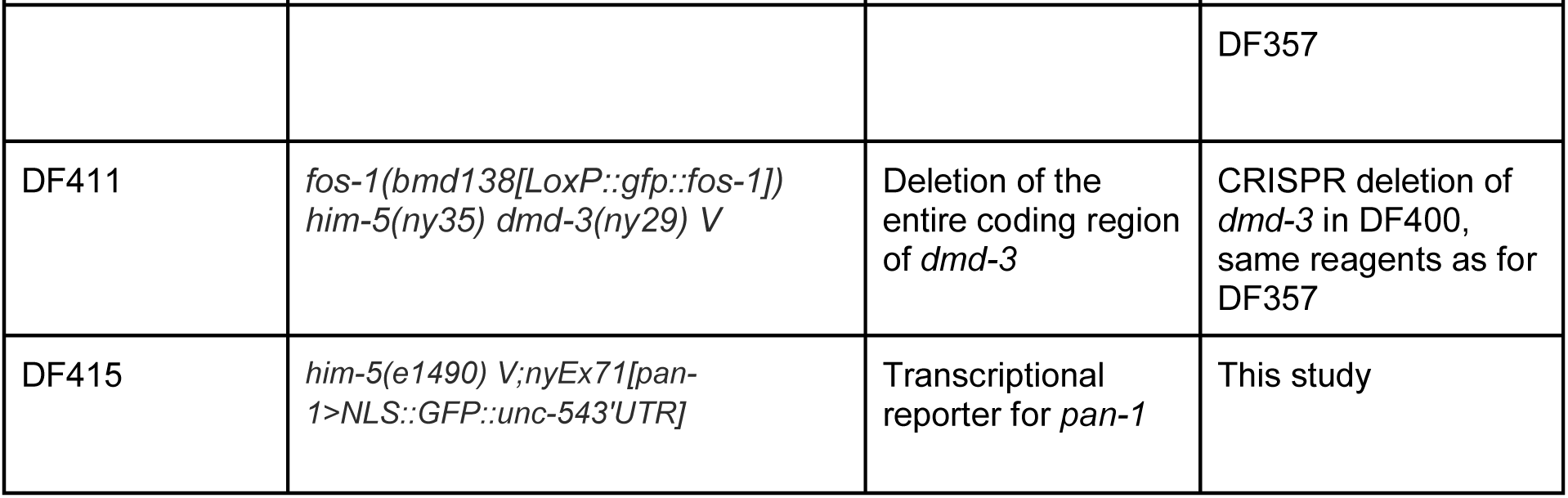
*C. elegans* strains used in this study (* We use the symbol “>” to designate that the promoter of a gene specified to the left of the symbol is used to drive expression of a marker construct to the right of the symbol.)

### Enriching males for ChIP-seq

To enrich worm collections for L4 stage males, we took advantage of a temperature-sensitive allele of the dosage compensation complex gene *dpy-28*(*y1*ts,mat) that causes specific hermaphrodite lethality at the restrictive temperature (Haque et al. 2024) (Fig. S1). First, worms were synchronized via bleaching and L1 arrest (Lewis and Fleming 1995). The arrested L1s were allowed to grow at the permissive temperature (15 °C) on 100 mm diameter NGM plates supplied with concentrated *E. coli* HB101, until these were packed with gravid hermaphrodites. Hermaphrodites were then collected into M9 buffer and bleached to release their embryos, which were incubated without food at the restrictive temperatures of 20 °C or 25 °C overnight. Only male L1s survived, which were placed on food and allowed to develop until GFP::DMD-3 was expressed in all tail tip nuclei (early L4, ∼26 hours at 25 °C or ∼34 hours at 20 °C). GFP::DMD-3 expression was confirmed by fluorescence microscopy. Sucrose floatation (Zanin et al. 2011) was used to clean the worms from bacteria and debris. Approximately 400 µl of packed L4 males per sample could be collected with this method. Samples were cross-linked by adding formaldehyde (2%) and incubating on a rocker at room temperature for 30 minutes. Afterward, glycine was added to a final concentration of 125 mM and incubated at room temperature for 5 minutes. Worm collections were washed with M9 buffer, and an aliquot was collected to confirm GFP::DMD-3 expression. Fixed worms were transferred to a 1-ml microcentrifuge tube containing 1x PBS and protease inhibitors, washed twice in this solution, the supernatant was removed and the samples either flash frozen in liquid nitrogen and stored at -80 °C, or immediately subjected to the ChIP-seq protocol.

**Figure S1.**
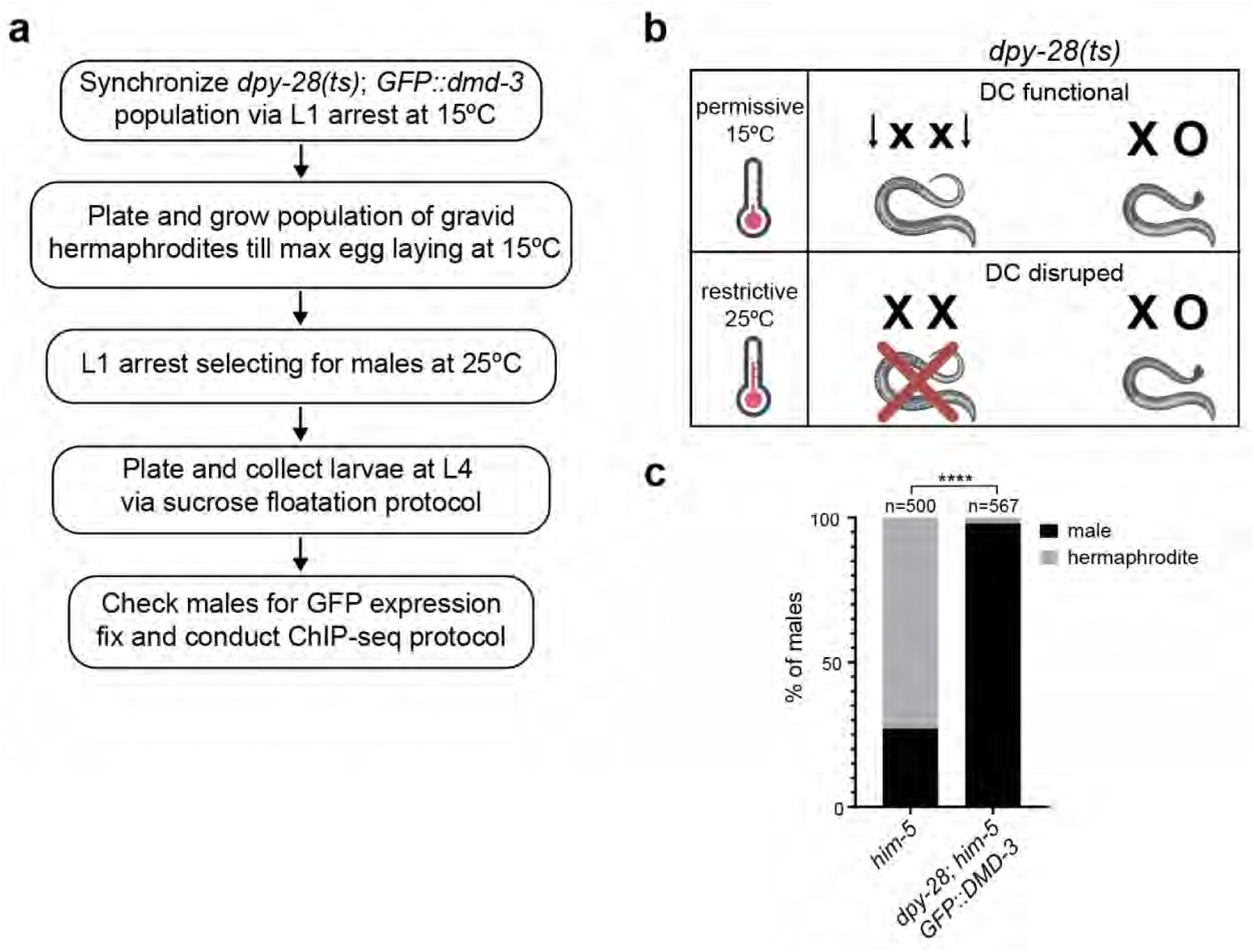
Strategy to enrich for L4 males with a temperature-sensitive mutation of *dpy-28.* a) Workflow for obtaining synchronized GFP::DMD-3-expressing L4 males for ChIP-seq. DMD-3::GFP positive males were selected by synchronizing worms and shifting from permissive to restrictive temperatures for the *dpy-28(*ts) allele. b) Schematic depicting the outcome of growing *dpy-28(ts)* worms at low and high temperatures. At the restrictive temperature, hermaphrodite development arrests during embryogenesis, while male development continues. c) When grown at 25 °C, the frequency of males in the *dpy-28(ts); him-5 GFP::dmd-3* strain DF301 reaches 98% compared to 27% in *him-5* control strain CB4088 (****p=<0.0001, Fisher’s exact test). The figure was created with the help of BioRender.com.

### ChIP-seq and NGS library preparation

ChIP-seq experiments were performed on collected males using the protocol as described (Albritton et al. 2017; Askjaer et al. 2018). Briefly, samples were cross-linked by adding formaldehyde (2%) and incubating on a rocker at room temperature for 30 minutes. Glycine was added to a final concentration of 125mM and incubated at room temperature for 5 minutes. For ChIP extract preparation, samples were dounce homogenized in FA-buffer (50 mM HEPES/KOH pH 7.5, 1 mM EDTA, 1% Triton X-100, 0.1% sodium deoxycholate; 150 mM NaCl). Sarkosyl (0.1% sodium lauroyl sarcosinate) was added to samples before sonication. Samples were sonicated to enrich chromatin fragments between 200bp and 800bp in length, and protein concentration was determined via Bradford Assay (Askjaer et al. 2018). For chromatin immunoprecipitation, 4 mg of ChIP extract was treated with 8 µg GFP antibody ab290 (Abcam) per ChIP (Ercan et al. 2007).

ChIP-DNA samples were ligated to barcode adaptors from the NEXTflex ChIP-seq Barcodes kit (Bioo Scientific) using the NEXTflex ChIP-seq kit (Bioo Scientific) following the manufacturer’s protocol with the following alterations: the initial size selection step (step B1) was skipped, the number of PCR cycles in the amplification step was 15. After amplification, 200-600bp fragments were size-selected using SPRIselect beads (Beckman Coulter).

Single-end 150bp sequencing (for three DMD-3 ChIP-seq libraries and their three corresponding no-antibody inputs, 6 libraries total) was performed using the Illumina Nextseq 500 platform at the New York University Center for Genomics and Systems Biology New York, NY. All sequencing data are available at Gene Expression Omnibus under GEO Accession # GSE282566

### ChIP-seq data processing

Adapter sequences from raw reads were trimmed and filtered using the BBduk shell script from BBmap Version 38.42 (Bushnell et al. 2017). We aligned 150bp single-end ChIP-seq reads to *C. elegans* genome version WBcel235/ce11 using bowtie2 version 2.3.2 under the default parameters (Langmead and Salzberg 2012). Duplicate reads were removed using MAC2 version 2.1.1 (Zhang et al. 2008), and BAM files were sorted and indexed using Samtools version 1.6 (Ramirez-Gonzalez et al. 2012). DeepTools version 3.3.1 was employed for coverage generations and normalization by subtracting by input via bamCompare settings: CPM, bin-size of 10bp, ignore duplicates, extend reads to 391-400bp, exclude chrM, minMappingQuality 20 and to remove blacklisted regions (Ramírez et al. 2016; Amemiya et al. 2019). Using MACS2, we predicted the estimated fragment size and called peak enrichment regions. Peaks were called on a merged BAM file using a threshold false discovery rate (FDR) of 0.05. All scripts used to process the data are available on Git Hub (https://github.com/classymagpie/DMD3_ChIPseq).

### *De novo* motif analysis of DMD-3 ChIP-seq enrichment peaks

Using default settings, MEME-ChIP was used to scan 100-bp regions centered around DMD-3 ChIP-seq peak summits. The background sequence generation employs a Markov 2nd-order model from shuffled primary sequences as input. Also, to help identify which motifs are centrally enriched, we used another MEME Suite tool, CentriMo. For CentriMo analysis (as well as FIMO below), we extended peak regions to 500bp centered on peak summits as recommended (Machanick and Bailey 2011). The parameters used for MEME-ChIP/CentriMo are as follows:

> “meme-chip -oc . -time 240 -ccut 100 -fdesc description -dna -order 2 -minw 6 - maxw 15 -db db/WORM/uniprobe_worm.meme -meme-mod zoops -meme-nmotifs 7 -meme-searchsize 100000 -streme-pvt 0.05 -streme-totallength 4000000 -centrimo-score 5.0 -centrimo-ethresh 10.0 target.fa”

To Identify the number of DMD-3 motif occurrences within DMD-3 enrichment peaks, we utilized the MEME Suite tool FIMO, which was run using default settings and a p-value threshold of 0.001 (Grant et al. 2011). As an alternative *de novo* motif discovery tool, we scanned enrichment peaks 500bp centered on peak summits using the HOMER tool Perl script findMotifsGenome.pl using default settings (Heinz et al. 2010).

We retrieved from the GTRD database 9394 EOR-1 ChIP-seq binding regions derived from three experiments from the modENCODE project (Kolmykov et al. 2021). Then, we applied the bedtools intersect function (bedtools version 2.29.2)

> “bedtools intersect -a DMD3peaksINbed.bed -b EOR1_peaksINbed.bed -u > DMD3_EOR1_OLpeaks_U_output.txt”

This determined how many of our DMD-3 ChIP-seq peaks (1755) intersected with those called for EOR-1 (9394). We then determined target genes within a 10-kb window around TSS (6217 total genes for EOR-1 vs 6061 genes for DMD-3).

### Transgenes and Constructs

To investigate the potential DMD-3 binding motif, we designed a 35bp oligo with the motif sequence identified by MEME-ChIP in the center (15bp, uppercase) flanked by 10bp on each side derived from a motif region upstream of *pop-1* (I: 2831554..2831588; ctacaccaatTTCTCTGCGTCTCTCgctttcatcg). As a negative control, we scrambled the central motif sequence to ctacaccaatCTGGTTTTTCCCTCCgctttcatcg. We ordered these oligos and their reverse complements from IDT. To make dsDNA, each complementary set of oligos was combined at equimolar concentrations and subjected to the following melting and re-annealing protocol: 95 °C for 2 minutes; 85 °C for 10 seconds, 75 °C for 10 seconds, 65 °C for 10 seconds, 55 °C for 1 minute, 45 °C for 30 seconds, 35 °C for 10 seconds, 25 °C for 10 seconds, 4 °C hold. Each dsDNA was injected into CB4088 hermaphrodites at a final concentration of 10 µM with 9 ng plasmid pCFJ90 (Addgene #19327, *myo-2>mCherry::unc-54 ‘UTR*) as an injection marker. Transgenic F1 male progeny were screened for TTM defects.

A transcriptional reporter for *pan-1,* driving 4xNLS::GFP with 2427nt of the regulatory region of *pan-1*, including the first exon and intron (III: 4560495..4562921), was made by overlap PCR of two PCR products (Nelson and Fitch 2011). The 5’ piece was made with primers pan1_TR_F1+MosTI_IV8.6:

*TTTAAATACTGGCTTAAAATGTTACGGGTCCAGGT*CGTCATTGCCAACGAACTTC, containing an overlap with a MosSci insertion site (italics) for alternative use, and pan1_TR_R3+NLS:

*GTCCTCCTGAAAATGTTCTATGTTATGTTAGTATC*TTCTTGGATATTATCAAACTCCG T with an overhang in plasmid pPD122.13 (in italics), from *C. elegans* N2 genomic DNA. The GFP-containing piece was made with primers pPD122.13_F:

GATACTAACATAACATAGAACATTTTCA and pPD122.13_R2: GTTCAGATGAGAGGAGCG from plasmid pPD122.13 (4xNLS-GFP::let-858 Addgene #19641) For both, PrimeStar Max DNA polymerase (Takara) was used with 35 cycles, annealing temperature 55 °C, extension time 15 seconds. The PCR products were purified (Promega Wizard SV Gel and PCR clean-up system), annealed for 13 cycles (Prime Star Max, annealing temperature 55 °C, extension time 15 seconds), and the product used for final extension with primers Pan1TR_F2: ACTTCTCGAACACGGAACAG and pPD122.13_R1*: AATGTTTAGATTTGGATTGA (35 cycles, same PCR conditions as above). The final product was purified and micro-injected into CB4088 hermaphrodites at a concentration of approximately 1 fmol with myo-2>mCherry (plasmid pCFJ90) as injection marker. Micro-injections were done as previously described (Evans 2006).

### CRISPR Cas9 genome editing

CRISPR Cas9 genome editing using guide-RNA-loaded Cas9 protein was used to insert GFP into the *pan-1* locus, to delete peak regions, and to introduce the *him-5(-)* and *dmd-3(-)* mutations as in (Kiontke, Fernandez, et al. 2024). Sequences of all guide RNAs, homology repair templates (HRTs), and primers are found in Table 2. For some experiments, the *dpy-10* co-CRISPR strategy was used (Arribere et al. 2014). All guide RNAs were designed with the tools in Benchling (https://www.benchling.com). RNAs and the Cas9 enzyme were ordered from IDT. For deletion of larger pieces, two guide RNAs and a 70 nt long HRT bridging the gap were used. Edits were identified by PCR with primers flanking the gap or the insertion except for *him-5(-)* and *dmd-3(-)*, where we only confirmed the phenotype (many males in the progeny of hermaphrodites or males with tail tip defects, respectively). Cas9-guideRNA complexes were prepared following the protocol provided by IDT. Briefly, crRNAs and tracer RNAs were prepared in IDT duplex buffer to a concentration of 200 µM. 0.5 µl of each were annealed at 95 °C for 5 minutes. The guide RNA was diluted with water or with the dpy-10 guide RNA (10 µM) to 62 µM. 0.5 µl of this RNA mix was combined with 0.5 µl Cas9 protein and incubated for 5 minutes at room temperature. For the injection mix, the RNP complex was combined with the respective HRT for the targeted gene at a final concentration of 20 µM for dsDNA and 2 µM for ssDNA in a volume of 10 µl. For co-CRISPR, the dpy-10 HTR was added at a final concentration of 0.1 µM. The HRT for attaching GFP at the 3’ end of *pan-1* was made by PCR (Takara PrimeStar Max) from plasmid pPD95.75 (Addene #1494) that contains GFP. The product was purified (Promega Wizard SV Gel and PCR clean-up system) and its concentration adjusted to 100 ng/µl (measured by Nanodrop Thermo Scientific). Before assembling the injection mix, this double-stranded DNA was heated to 95 °C for 5 minutes and put on ice. The HRT for all other edits was ordered from IDT as single-stranded oligonucleotides. The mix was injected into both gonads of 8 to 10 young hermaphrodites. One to five L4 progeny of the injected worms were picked onto individual plates and allowed to lay eggs for one day after which they were picked into lysis buffer (Fay and Bender 2006) containing 10 µM proteinase K, frozen in liquid nitrogen and thawed 3 times and incubated at 60 °C for 90 minutes followed by enzyme inactivation at 95 °C for 15 minutes. This lysis was used for PCR with primers flanking the HRT insertion site (see Table 2). After a line with an edit was identified, this process was repeated until the edit was homozygous.

**Table 2.**
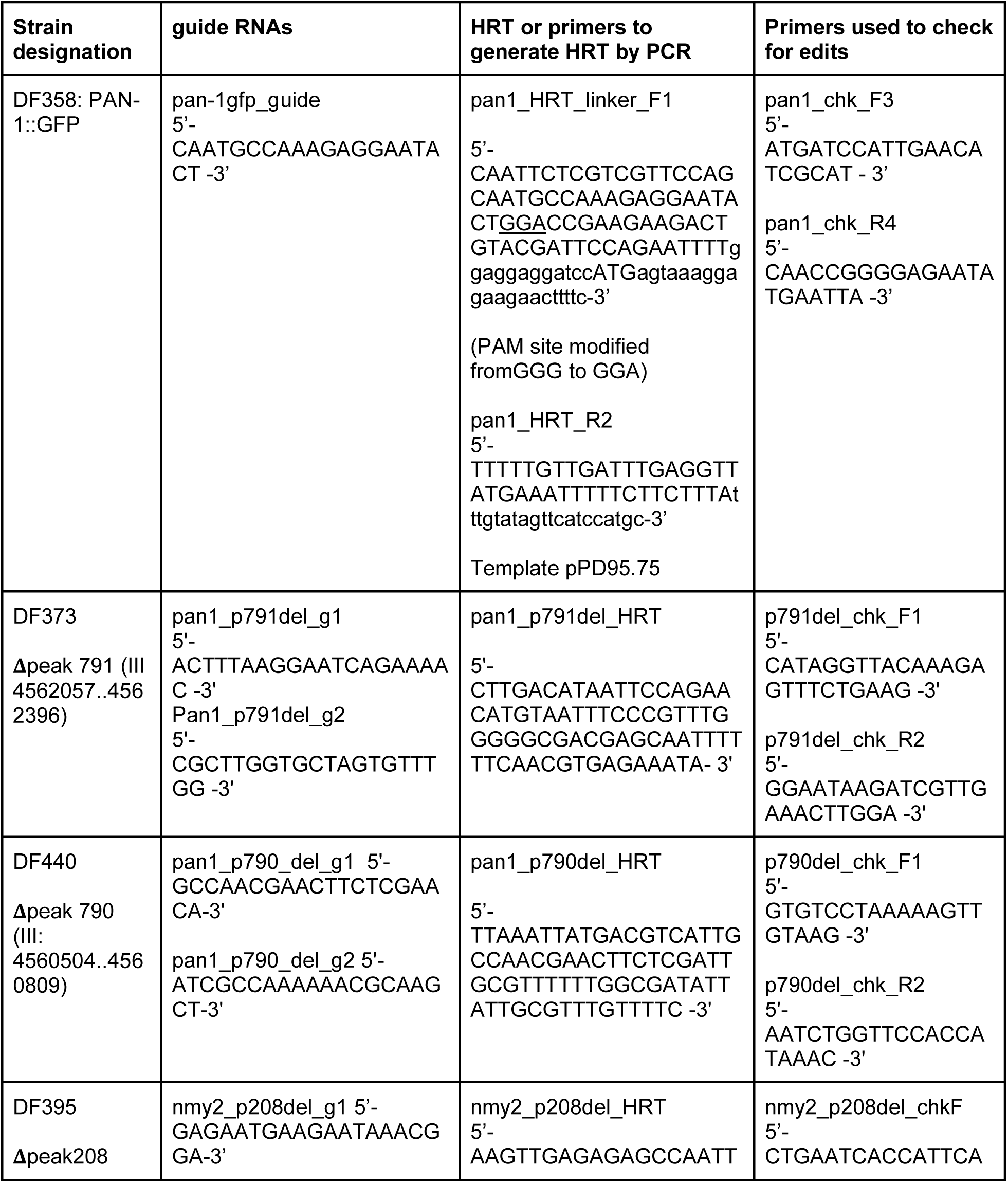
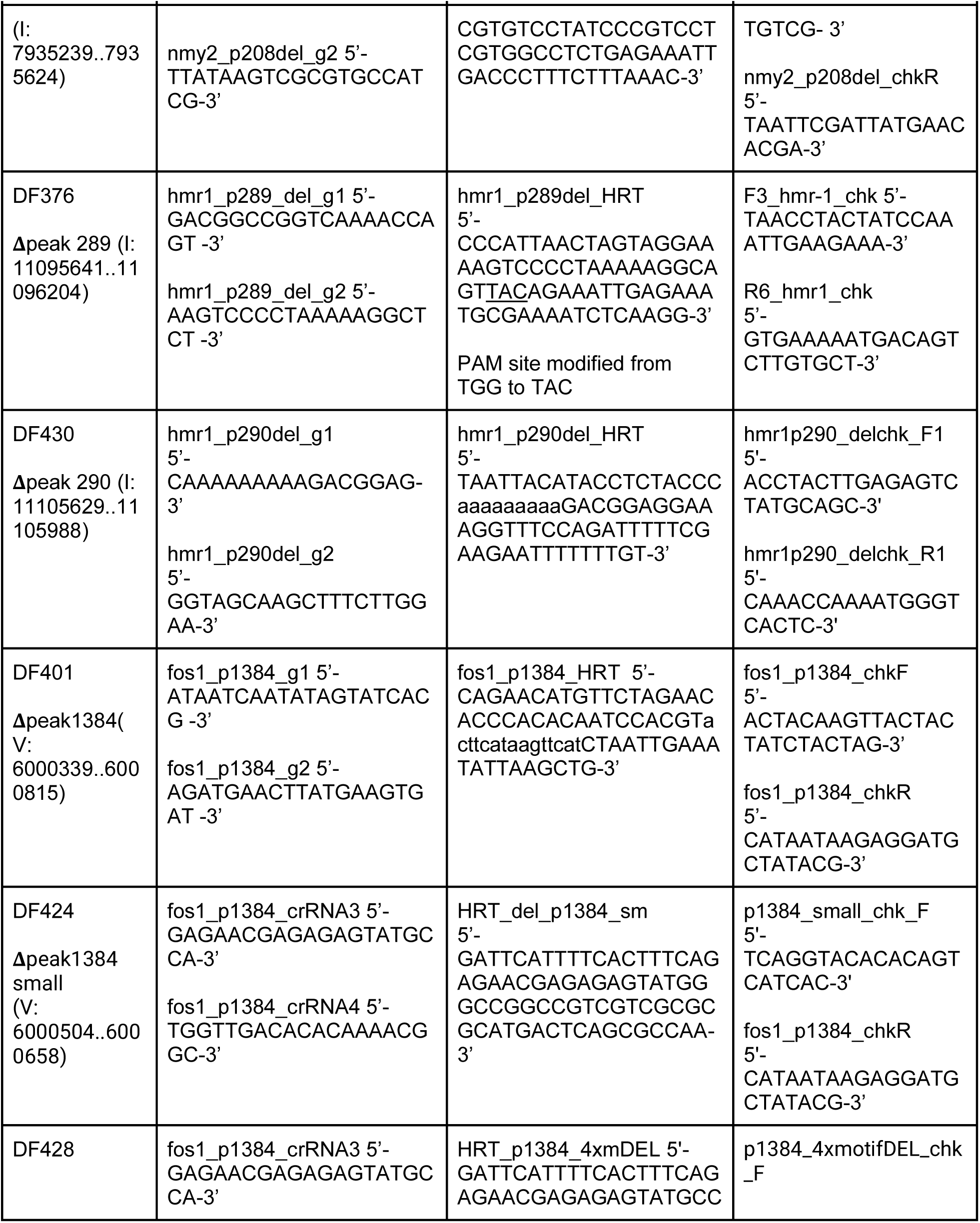
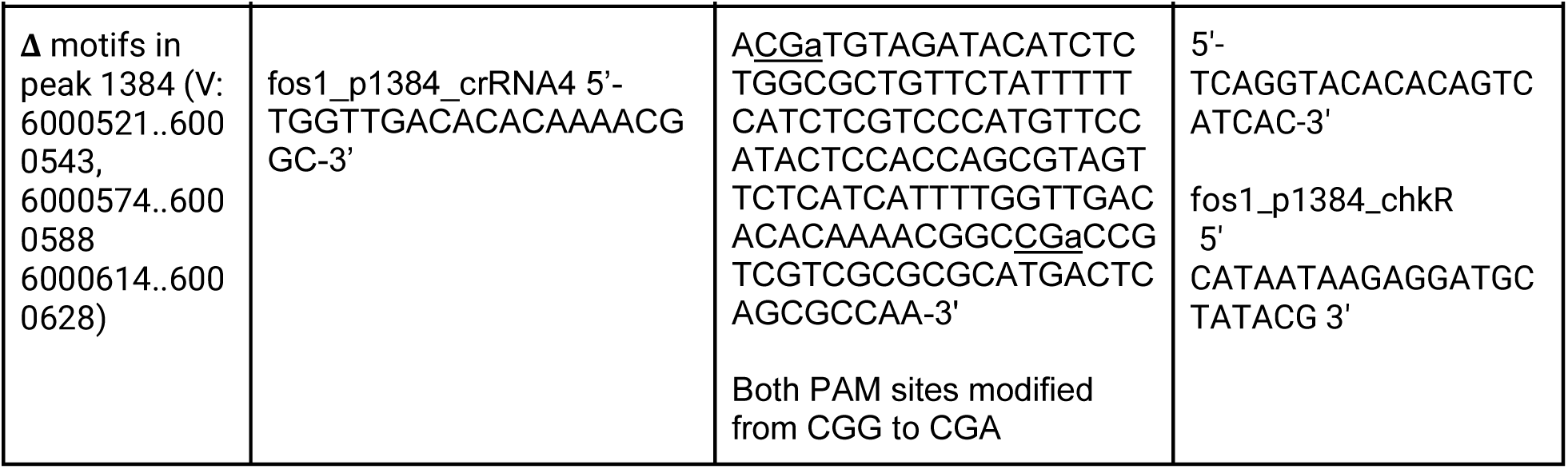
Strains made by genome editing and CRISPR reagents.

### Microscopy and imaging

Worms were immobilized in a 20 mM sodium azide solution in M9 buffer and mounted on a pad of 4% Noble agar in water supplemented with 20 mM sodium azide. Worms were then visualized using a Zeiss AxioImager with Colibri LED illumination and an Apotome. Z-stacks were taken with at most 0.5 µm distance between slices. If signal intensity was too low for Apotome imaging, we employed the ZenBlue (2.0) deconvolution module using the ‘good/fast iterative’ setting if image quality needed to be improved.

### Fluorescence signal quantification

To quantify the intensity of the NMY-2::mKate fluorescence in the tail tip of males with and without enrichment peak deletions, we synchronized populations of DF343 (NMY-2::mKate + Lifeact::GFP) and DF395 (Δ peak208 in NMY-2::mKate), and mounted L4 males of both strains onto the same slide, where they could be distinguished by GFP fluorescence. Image Z-stacks of tail tips were taken with 1s exposure time with a light source intensity of 25% or 30% for the red (mCherry) channel, only images with identical exposure and intensity settings were compared. The fluorescence signal intensity was determined using ZenBlue(2). We recorded the mean fluorescence intensity in a circle (17.339 µm2) centered on a region in the tail tip of similarly staged L4 males and in a region outside the animals for background subtraction. Plots were generated, and statistical analysis (unpaired t-test) was performed using GraphPad Prism version 10.0.0 for Mac OS, GraphPad Software, Boston, Massachusetts USA, www.graphpad.com.

## RESULTS

### Whole-worm L4 male-specific ChIP-seq identifies 1,755 DMD-3-associated enrichment peaks

To identify DMD-3 target sites in the *C. elegans* genome, we performed ChIP-seq using a strain (DF288) in which the endogenous DMD-3 protein is tagged N-terminally with GFP (Kiontke, Fernandez, et al. 2024) and an anti-GFP antibody to pull down DNA-bound DMD-3. We performed ChIP-seq in early L4 samples enriched for males. To this end, the strain contained the temperature-sensitive mutation of the dosage compensation complex gene *dpy-28*; in which approximately 98% of the population consisted of males at the restrictive temperature (25 °C) (Fig. S1). Males were collected 26 hours after L1 arrest at a time point corresponding to L4 larval substages L4.2-L4.3 when GFP::DMD-3 expression peaks in tail tip cells and TTM begins (Kiontke, Fernandez, et al. 2024). This is the same time point at which tail tips were collected by laser microdissection for differential expression (DE) RNA-seq analysis in a previous study (Kiontke, Herrera, et al. 2024).

Using MACS2 (Zhang et al. 2008), we identified 1,755 ChIP-seq enrichment peaks (FPR = 0.05, *d* = 290bp) (File S3, Fig. S2). As expected for legitimate transcription factor binding sites (Niu et al. 2011), 82% of the reads in the enrichment peaks map within 2kb upstream of transcription start sites (TSSs) in the *C. elegans* genome as annotated in WormBase (Davis et al. 2022). Of these, 69% map within 1kb upstream of the TSSs (Fig. 2). Another 12% fall into intergenic regions more than 1kb from a TSS. The remaining reads fall inside exons or introns (6%) or within 300bp from the 3’ end of gene (transcript) annotations (0.2%) (Fig. 2). We assigned enrichment peaks to genes using 4-kb, 6-kb, 8-kb, and 10-kb windows centered on the TSS. This resulted in 3636, 4437, 5228, or 6061 total genes represented in each window, respectively (Fig 2, File S3.).

**Figure S2.**
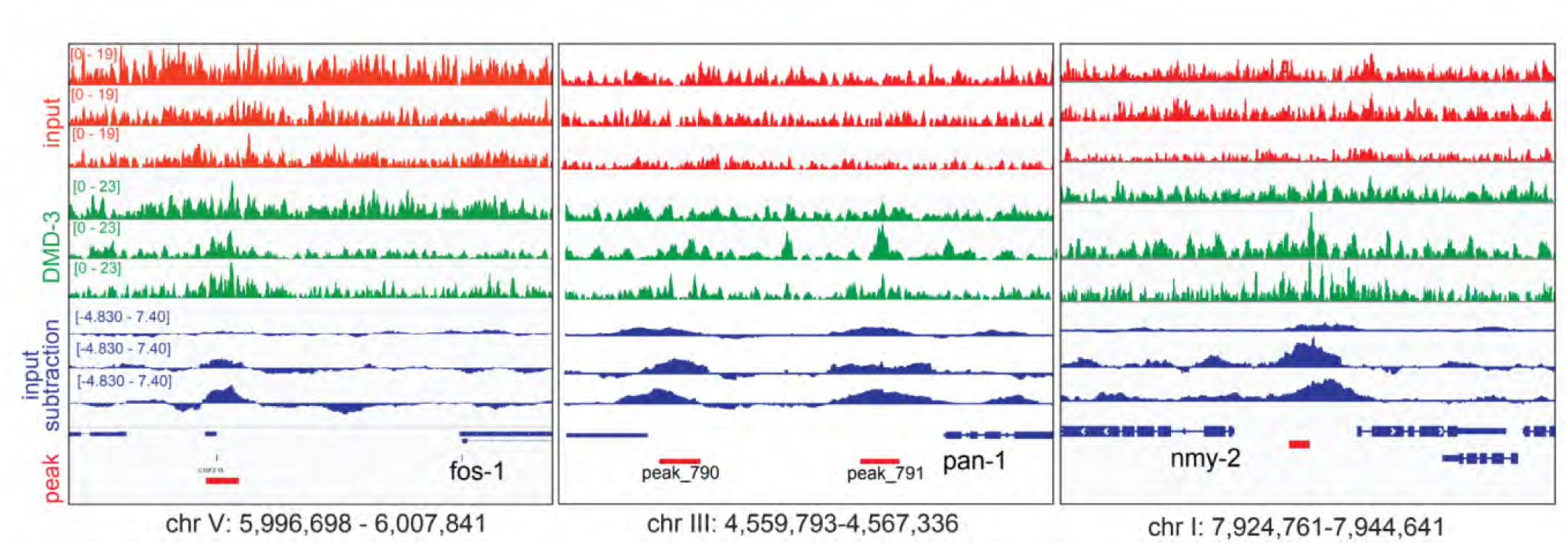
Sample DMD-3 ChIP-seq IGV coverage tracks covering *fos-1*, *pan-1* and *nmy-2* enhancer regions. The top three tracks (red) are input samples (no antibody treatment). The middle three tracks (green) are the DMD-3 pulldown samples. The lower three tracks (blue) represent ChIP coverage after subtracting the input. Below these tracks are the reference genome (genes represented by blocks and lines) and the peaks called using MACS2 (red blocks). Genes are noted under the genes, and the genomic location coordinates are denoted at the bottom.

**Figure 2.**
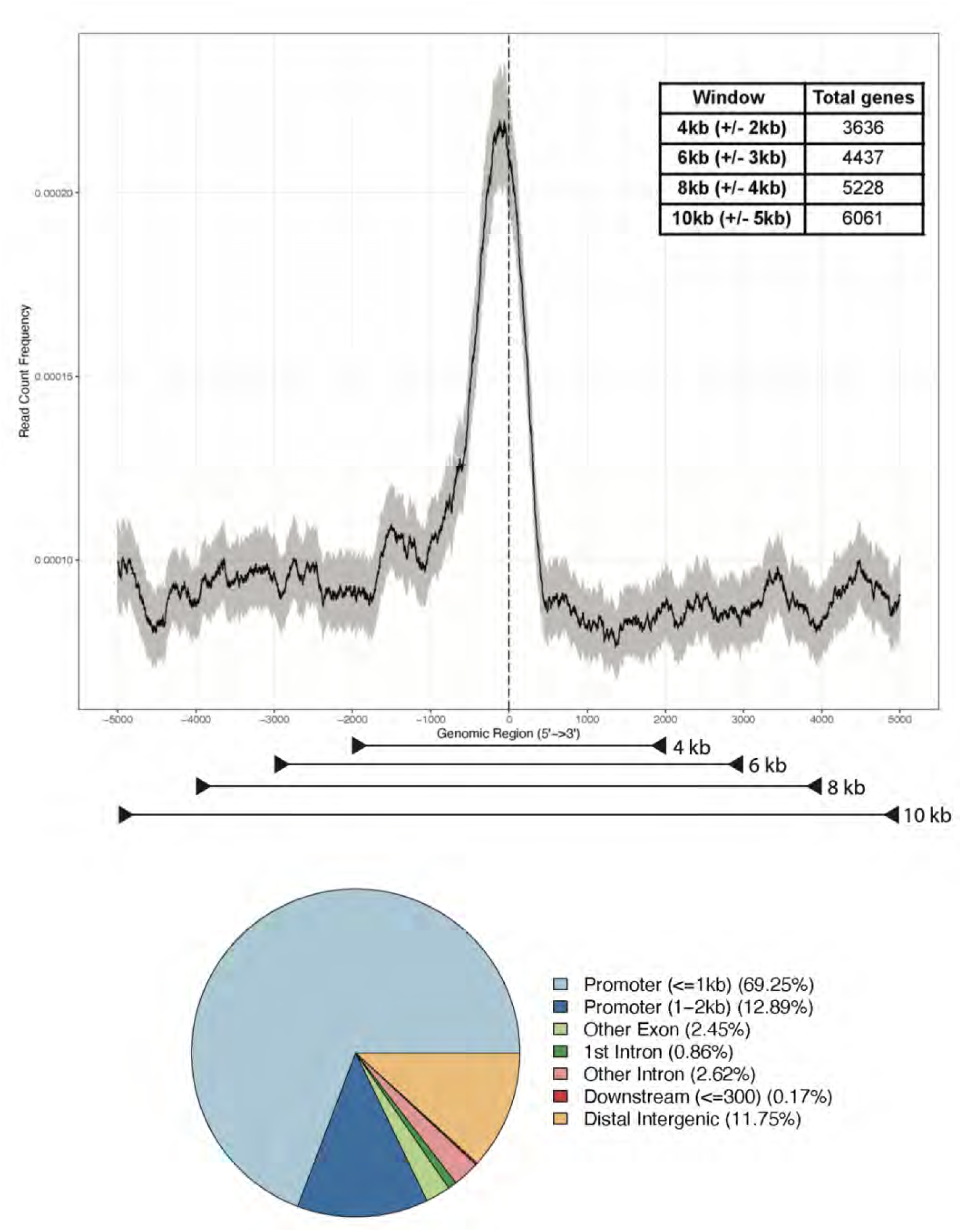
Profile of genomic reads corresponding to DMD-3 ChIP enrichment. a) Distribution of aligned reads of DMD-3 ChIP-seq peak regions plotted against genomic location in a 10-kb window centered on the transcription start sites of genes. The gray area around the black line represents the 95% confidence interval. The genomic windows used to assign genes to peaks are indicated below, gene numbers per window in the inset. b) Pie graph representing the percentage of aligned reads corresponding to DMD-3 ChIP-seq peak regions characterized by genomic annotation and distance to TSS(s).

### Identification of a DMD-3-associated consensus binding motif which is functionally important for TTM

To identify a putative DMD-3 recognition sequence, we scanned the 1,755 enrichment peak sequences for a shared motif using MEME-ChIP (Machanick and Bailey 2011). The predominant bases in the MEME-ChIP motif with the highest confidence score (Eval = 3.8e-177) are YTSTCTSYGTCTCYH (Fig. 3, File S1). According to the MEME-Suite tool FIMO (Grant et al. 2011), this motif is present at least once in 79% of the enrichment peaks (File S1, File S3). The alternative algorithm, HOMER (Heinz et al. 2010), identified a 12 nt motif nearly identical to the 15 nt MEME-ChIP motif and nested within it (TGTSTGTGTCTC Fig. 3). This motif is strikingly similar to the motif for the transcription factor EOR-1 (TCTCGCGTCTCT, Fig. 3) (Fornes et al. 2020). To confirm this overlap, we compared our DMD-3 ChIP-seq results against EOR-1 ChIP-seq data for L3 hermaphrodite and mixed-sex samples found in the Gene Transcription Regulation Database (GTRD) (Yevshin et al. 2019). Our analysis determined an EOR-1/DMD-3 overlap of 42% for peaks and 41% for target genes. This overlap includes peaks that do not contain the DMD-3 associated motif.

**Figure 3.**
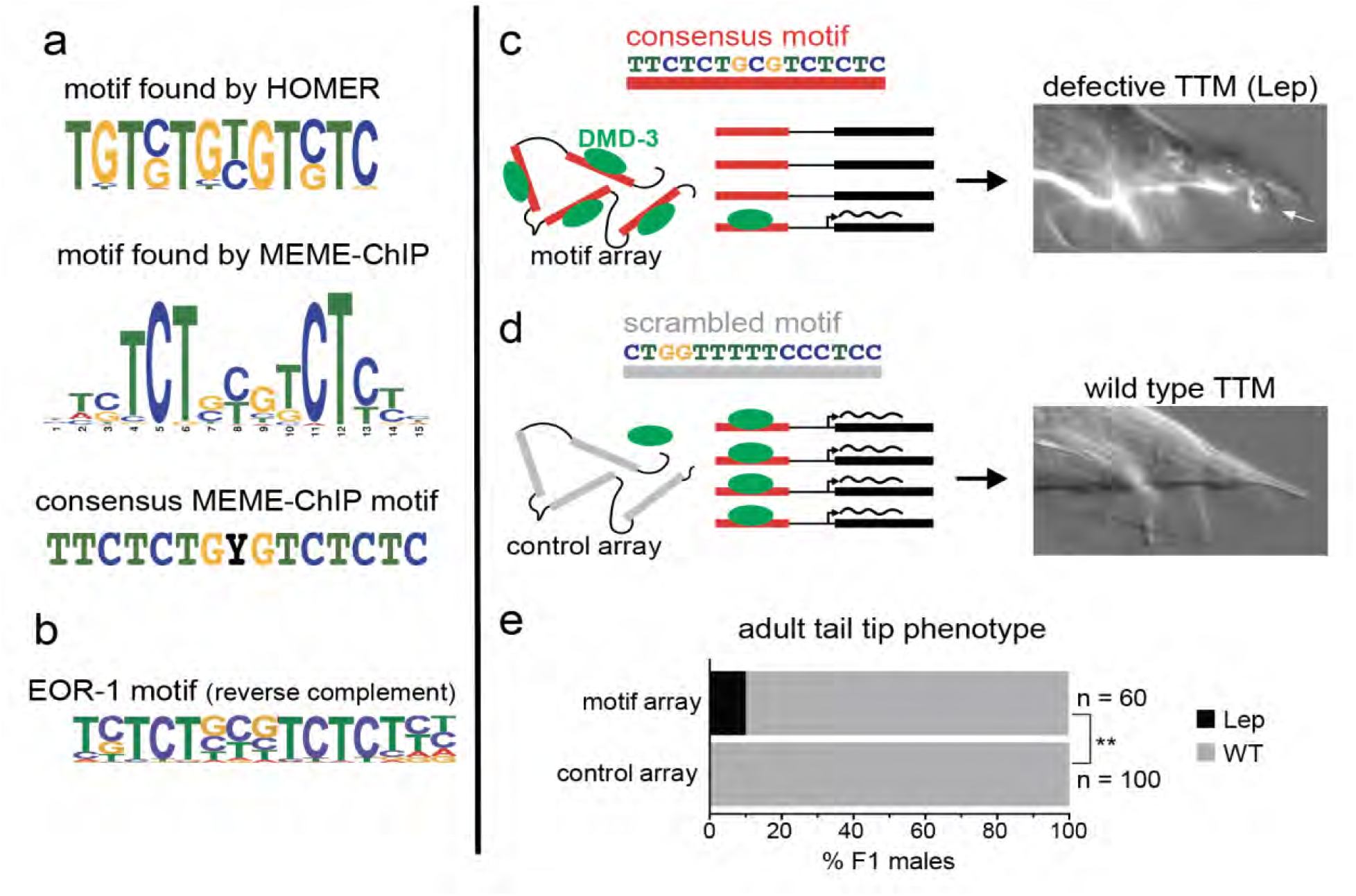
The DMD-3 associated motif determined via *de novo* motif discovery and array binding assay. a) *De novo* motifs in DMD-3 ChIP-seq enrichment peaks identified by HOMER and MEME-ChIP, and consensus motif used for validation experiments. b) Reverse complement of the EOR-1 motif (MA0543.2, JASPAR database (Fornes et al. 2020)). c) Schematic of the experimental design: a transgenic array with the 15bp motif consensus sequence binds to DMD-3 protein *in vivo,* sequestering it from its endogenous target genes and resulting in defective TTM (Lep adult tails). d) A control scrambled-motif array does not bind DMD-3 (WT adult tails). e) Quantification of results, (**p=0.0025, chi-square test).

To examine if the DMD-3-associated binding motif identified by MEME-ChIP is functional for TTM, we introduced an extrachromosomal array tiled with many copies of the predominant form of the motif (without ambiguities, TTCTCTGCGTCTCTC) into the worms and tested if this would result in defective TTM (Fig. 3). The rationale for this experiment was that if DMD-3 binds to the motif, DMD-3 would be sequestered by the array away from its normal chromosomal sites in target genes, thus phenocopying a reduced-function *dmd-3* mutant. Similar experiments were performed to identify the DNA sequences sufficient to recruit the dosage compensation complex in *C. elegans* (McDonel et al. 2006). Ten percent of adult males carrying the array had Lep tails (n = 60, Fig. 3), a typical result of defective or delayed TTM (Nelson et al. 2011), whereas none (n = 88) had tail tip defects when a control array with a scrambled motif was used instead (p = 0.0025, chi-square test). The phenocopy effect disappeared after the F1 generation, suggesting that the transgene was silenced in later generations, as is typical for many repetitive arrays in *C. elegans* (Leyva-Díaz et al. 2017). These results are consistent with the motif identified by us being a *bona fide* regulatory sequence required for wild-type TTM, to which DMD-3 binds directly or in concert with another TF such as EOR-1. We therefore refer to this motif as the “DMD-3 associated motif”.

### Candidate genes regulated directly by DMD-3 in TTM

At the collection time point (26 h after arrested L1s are plated), DMD-3 is expressed in many cells of the developing tail and in some cells of the somatic gonad in addition to the tail tip cells. Because our ChIP-seq experiment was performed on whole worms, the identified DMD-3-associated sites (“peaks”) may correspond to genes that function in any of these tissues. To find genes that could be direct targets of DMD-3 specifically in the tail tip during TTM, we took advantage of two previous datasets for genes involved in TTM: a whole-genome RNAi study (Nelson et al. 2011) and a tail tip-specific RNA-seq study (Kiontke, Herrera, et al. 2024).

We looked for an overlap (intersection) between the TTM genes identified in these studies and the genes assigned to DMD-3 enrichment peaks within a 10-kb window. We used this large window because, while most transcription factor binding sites in *C. elegans* are within 2kb of a TSS, some can be as distant as 10kb (Berkseth et al. 2013; Van Nostrand and Kim 2013; Araya et al. 2014; Daugherty et al. 2017). Although this large window likely identifies many false positives, we reasoned that only true positives would be found in the tail tip-specific datasets. For comparison with genes differentially expressed in the tail tip, we reanalyzed the dataset by Kiontke et al. (Kiontke, Herrera, et al. 2024) with an adjusted p-value threshold of 0.05 (using DEseq2 (Love et al. 2014)). This resulted in 1,154 DE genes for the comparison of wild-type with *dmd-3(-)* male tail tips (File S2). The intersection of this gene set with our 6,061 ChIP-seq candidate genes yielded 228 genes likely directly regulated by DMD-3 in TTM (Fig. 4, Table S1). Similarly, we interrogated the list of 211 genes giving TTM phenotypes in the genome-wide RNAi screen (Nelson et al. 2011). We found 48 DMD-3 targets among these genes (Fig. 4, Table S1). Considering both datasets, 270 genes are candidates for being both direct targets of DMD-3 and being involved in TTM (Fig. 4, Table S1). Although 42 genes are direct targets and yield an RNAi TTM phenotype, they were not differentially expressed in the RNA-seq analysis of tail tips of early L4 males (Kiontke, Herrera, et al. 2024). Regulation of these genes in the tail tip may occur at a different time during TTM or in a different tissue that affects TTM tail-tip cell non-autonomously.

**Figure 4.**
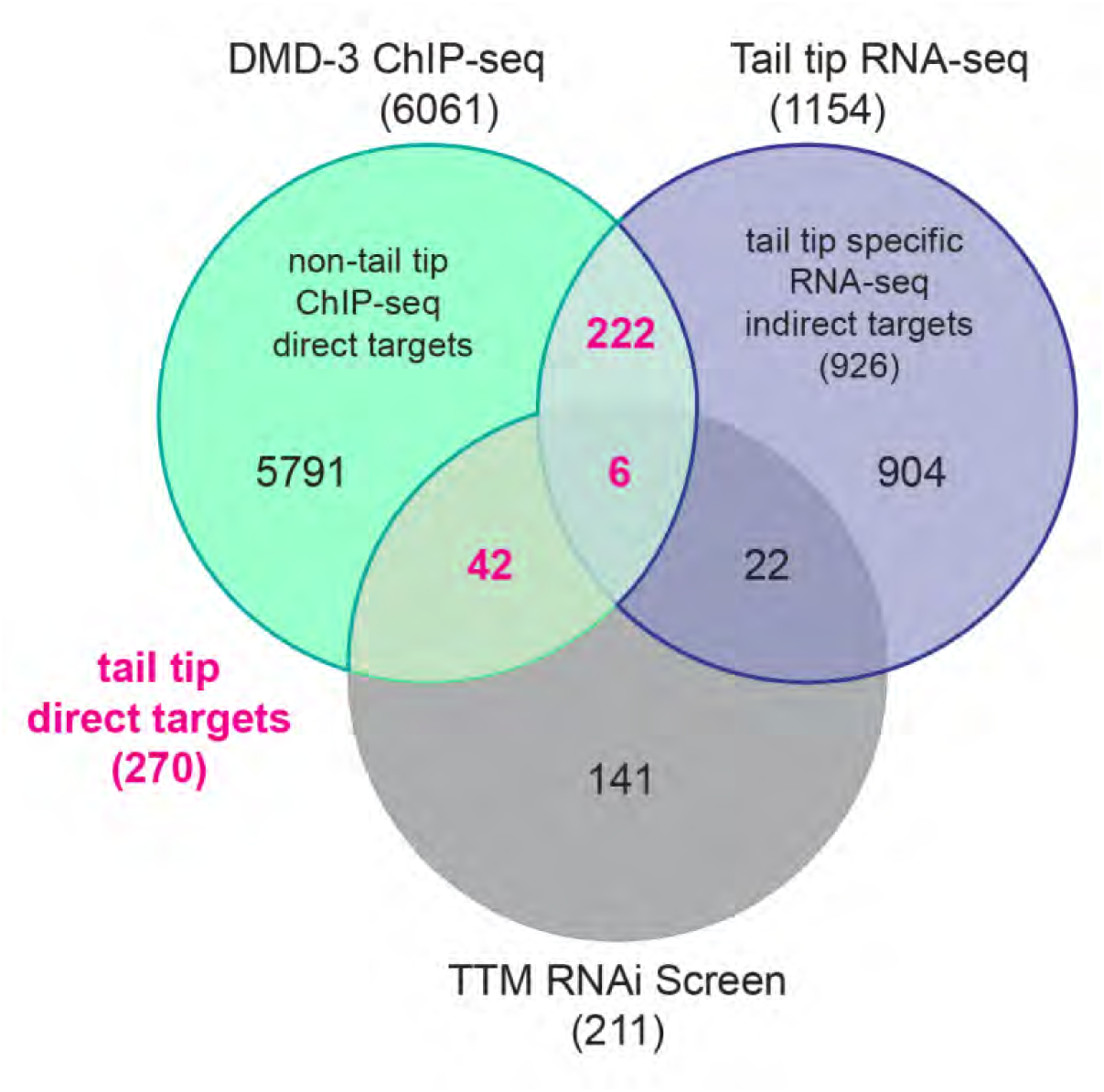
Overlap of TTM genes assigned to ChIP-seq peaks within a 10-kb window (green), DE genes in an RNA-seq analysis comparing tail tips isolated from WT *vs*. *dmd-3(lf)* males (Kiontke, Herrera, et al., 2024; blue), and a postembryonic RNAi screen for genes producing TTM phenotypes (Nelson et al., 2011; gray).

Mason et al. (2008) found evidence that *dmd-3* autoregulates and that this autoregulation depends on *mab-3*. For this reason, we wondered if *dmd-3* and *mab-3* are direct targets of DMD-3, although RNAi against these genes does not produce a phenotype and *mab-3* was not DE in the RNA-seq experiments. We did not find a DMD-3 enrichment peak in the cis-regulatory region of *dmd-3*. However, there were six DMD-3 enrichment peaks in an area 5kb around the TSS of *mab-3* with together 13 DMD-3 associated motifs. Thus, *mab-3* but not *dmd-3* is likely a direct target of DMD-3.

### Classification of TTM-relevant direct DMD-3 targets

To identify possible roles that the DMD-3 direct target genes play in TTM and to help select genes for further validation, we used gene descriptions in WormBase (Davis et al. 2022) to manually classify the 270 TTM-relevant DMD-3 targets into 17 broad primary functional categories (Table 3, Table S1, note that many genes can be assigned to several categories, here, we selected only one per gene). In our re-analysis of the RNA-seq data (Kiontke, Herrera, et al. 2024), we found 91 transcription factors (TFs) and 13 transcription cofactors downstream of DMD-3. Comparing this dataset and the TFs found in the RNAi-screen (Nelson et al. 2011) with our ChIP-seq shows that 28 TFs and 7 TF cofactors are direct targets of DMD-3. Of these regulators, two thirds are repressed by DMD-3 (Table 4, Table S1). Among the direct targets of DMD-3 are 24 kinases, phosphatases, GTPases and their regulators (classified previously as effectors (Kiontke, Herrera, et al. 2024)). DMD-3 direct targets further include genes that encode proteins with known roles in morphogenesis, including proteins of the cytoskeleton (e.g. NMY-2), of cell-cell junctions (e.g. AJM-1, INX-12 and INX-13), components of the WNT polarity pathway (SYS-1, LIT-1 and LIN-17) and proteins involved in vesicular trafficking (Table 5, Table S1). In addition, DMD-3 directly regulates 11 genes involved in the response to unfolded proteins (UPR) and ER stress, including for the heat-shock chaperones HSP-1, HSP-3 and HSP-90. Accordingly, a GO enrichment analysis (via WormBase (Angeles-Albores et al. 2016; Angeles-Albores et al. 2018)) finds as a highly enriched GO terms “DNA-binding transcription factor binding” (GO:0140297), “adherens junction” (GO:0005912) and “protein folding chaperone” (GO:0044183). Among the adherens junction genes is *hmr-1*, which we studied further below.

**Table 3.** Primary functional categories for 270 TTM-relevant direct DMD-3 targets. See Table S1 for all 270 genes with more detailed information.

| Functional category | number of direct targets |
| --- | --- |
| Cell cycle | 8 |
| Chromatin regulation | 8 |
| Endoplasmic reticulum | 9 |
| Extracellular matrix/cuticle | 26 |
| Host defense | 2 |
| Junctions | 7 |
| Membrane transport | 14 |
| Metabolic | 23 |
| Nuclear transport/Pore | 4 |
| Polarity/cytoskeleton | 12 |
| Post-transcriptional regulation | 22 |
| Protein folding | 11 |
| Signal transduction | 25 |
| Trafficking/endocytosis | 7 |
| Transcription factor | 28 |
| Transcription cofactor | 8 |
| Translational machinery | 3 |
| Unknown | 53 |

**Table 4.**
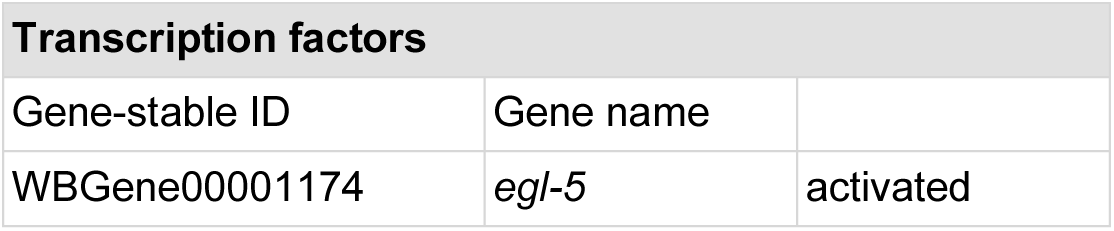
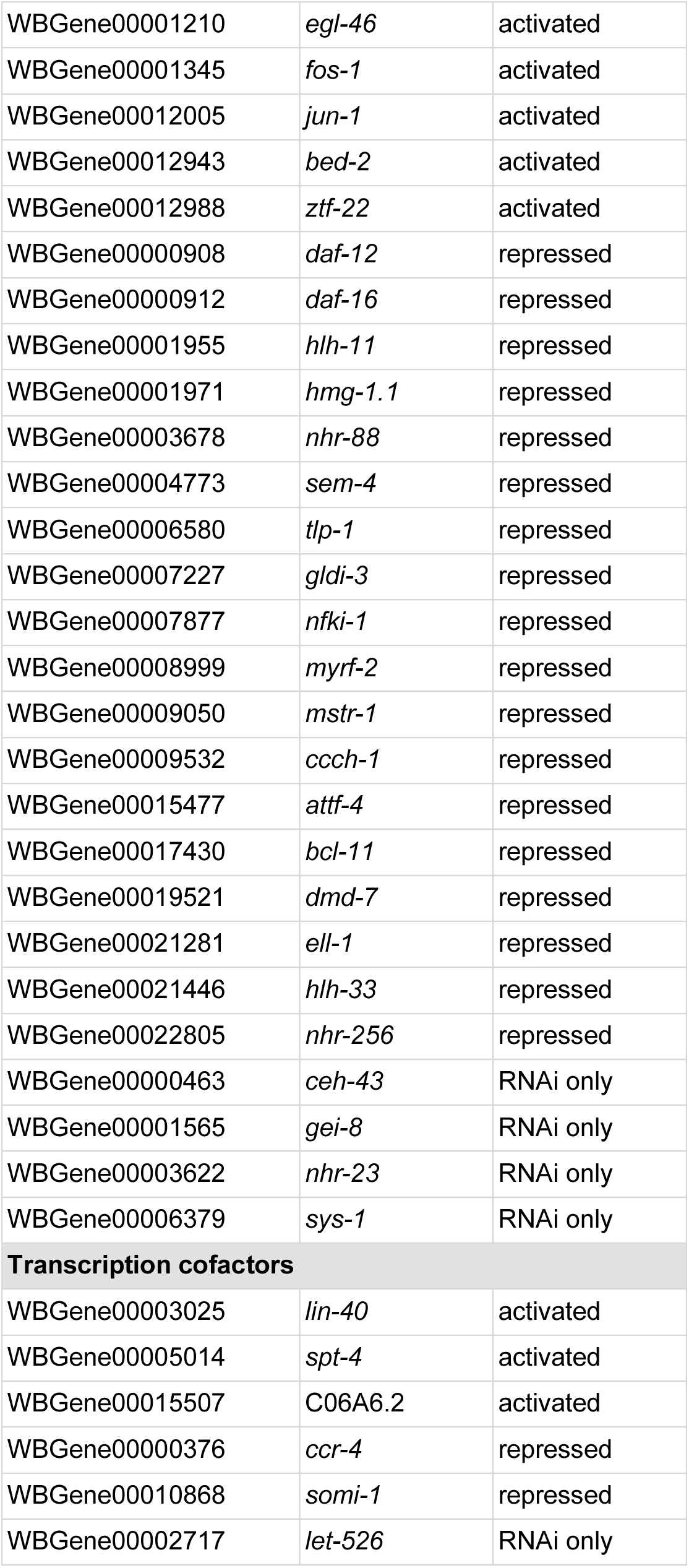
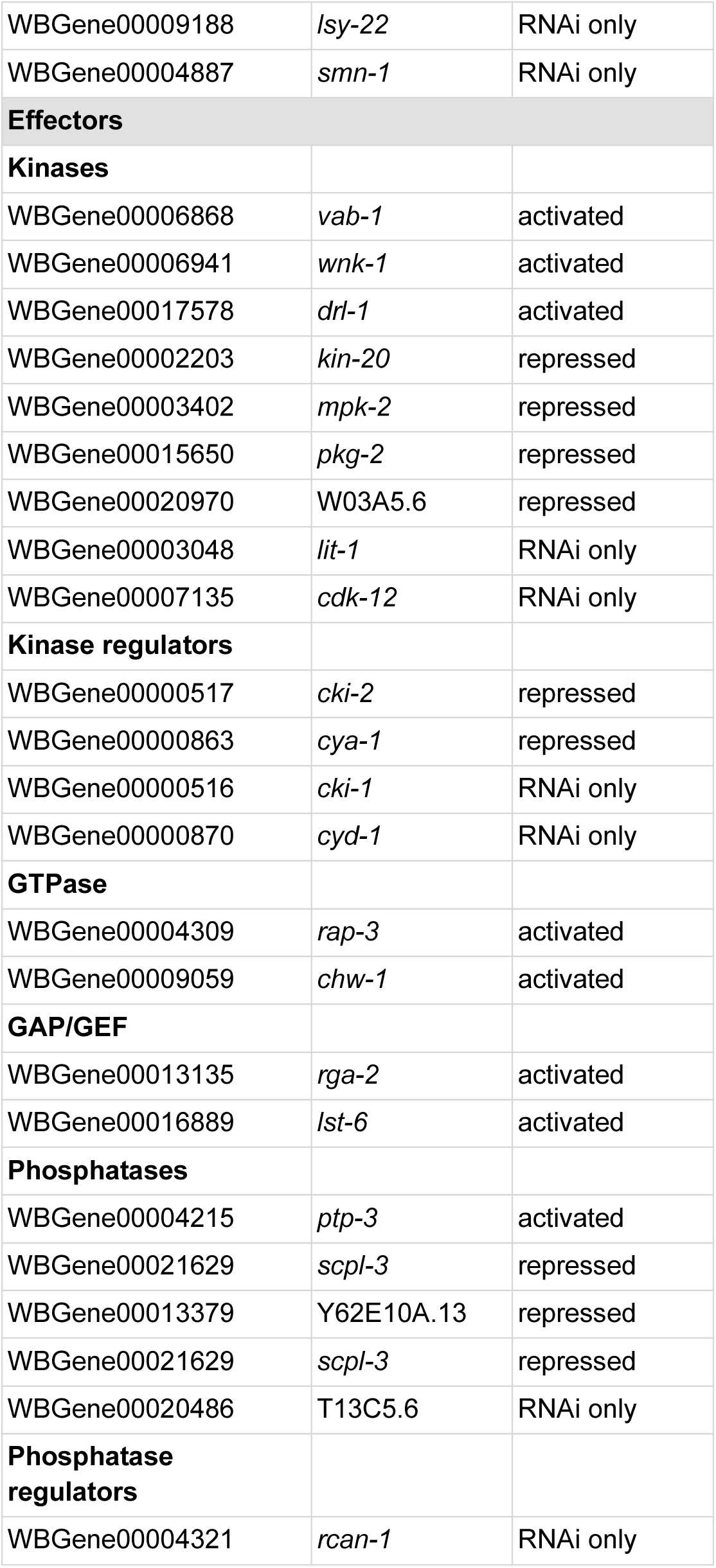
Transcription factors (TF) transcriptional cofactors (CF (Horowitz et al. 2023)) and “effectors” that are direct targets of DMD-3 in TTM. The third column indicates whether the gene has an RNAi phenotype (Nelson et al. 2011), or was activated or repressed by DMD-3 in the RNA-seq analysis (Kiontke, Herrera, et al. 2024).

**Table 5.** Universal and conditional module direct targets. The third column indicates whether the gene has an RNAi phenotype (Nelson et al. 2011), or was activated or repressed by DMD-3 in the RNA-seq analysis (Kiontke, Herrera, et al. 2024).

| **Direct targets within universal modules from Kiontke et al. (2024)** |  |  |  |
| --- | --- | --- | --- |
| **Cytoskeleton** |  |  |  |
| **Gene-stable ID** | **gene name** | **description/function** |  |
| WBGene00015687 | *chdp-1* | calponin homolog | activated |
| WBGene00000231 | *atx-2* | ataxin | activated |
| WBGene00019184 | H10E21.4 | predicted actin or actin-binding cytoskeletal protein | repressed |
| WBGene00006839 | *unc-115* | actin binding LIM protein family | repressed |
| WBGene00003777 | *nmy-2* | non-muscle myosin | RNAi |
| WBGene00002218 | *klp-6* | kinesin, cytoskeletal motor activity | RNAi |
| WBGene00002219 | *klp-7* | kinesin, microtubule binding | RNAi |
| WBGene00019128 | F59G1.4 | armadillo repeat containing protein, cilium assembly | RNAi |
| **Polarity** |  |  |  |
| WBGene00006769 | *unc-33* | establishment/maintenance of actin cytoskeleton polarity | activated |
| WBGene00008555 | F07C6.4 | establishment of epithelial cell polarity | activated |
| WBGene00003006 | *lin-17* | Wnt pathway, receptor | activated |
| WBGene00003048 | *lit-1* | Wnt pathway, kinase | RNAi |
| **Cell-matrix adhesion** |  |  |  |
| WBGene00001980 | *hmr-1* | cadherin | activated |
| WBGene00002135 | *inx-13* | innexin, gap junction | activated/RNAi |
| WBGene00021009 | *afd-1* | cell adhesion molecule binding | repressed |
| WBGene00003089 | *ltd-1* | cell-cell junction | repressed |
| WBGene00017931 | *picc-1* | adherens junction | repressed |
| WBGene00000100 | *ajm-1* | adherens junction | RNAi |
| WBGene00002134 | *inx-12* | innexin, gap junction | RNAi |

| Vesicular trafficking/endocytosis |  |  |  |
| --- | --- | --- | --- |
| WBGene00021292 | *copb-1* | vesicle-mediated transport | activated |
| WBGene00003589 | *nex-2* | annexin | activated |
| WBGene00006623 | *try-5* | secretory vesicle | activated |
| WBGene00011742 | *tgn-38* | trans-golgi network protein homolog | activated |
| WBGene00002980 | *lgg-1* | cytoplasmic vesicle | repressed |
| WBGene00004788 | *sft-4* | surfeit 4 homolog | repressed |
| WBGene00000894 | *dab-1* | clathrin-coated vesicle | repressed/RNAi |
| WBGene00006373 | *syx-5* | vesicle-mediated transport | RNAi |
| Unfolded protein response |  |  |  |
| WBGene00000567 | *cnx-1* | calnexin | activated |
| WBGene00000379 | *cct-4* | chaperonin, ER unfolded protein response | activated |
| WBGene00000915 | *hsp-90* | unfolded protein binding | activated |
| WBGene00001031 | *dnj-13* | DnaJ heat shock protein family | activated |
| WBGene00001099 | *dsc-4* | phospholipid transporter, ER unfolded protein response | activated |
| WBGene00002005 | *hsp-1* | protein refolding | activated |
| WBGene00002007 | *hsp-3* | ER unfolded protein response | activated |
| WBGene00003025 | *lin-40* | TF, mitochondrial unfolded protein response | activated |
| WBGene00011480 | *enpl-1* | heat shock protein 90 beta family | activated |
| WBGene00003962 | *pdi-1* | protein disulfide isomerase | activated |
| WBGene00003964 | *pdi-3* | protein disulfide isomerase | activated |
| WBGene00009331 | F32D8.7 | Involved in PERK-mediated unfolded protein response | repressed |
| Direct targets within conditional modules from Kiontke et al. (2024) |  |  |  |
| collagens upregulated during TTM |  |  |  |
| Gene-stable ID | gene name | description/function |  |
| WBGene00000623 | *col-46* | collagen | activated |
| WBGene00000716 | *col-143* | collagen | activated |
| WBGene00018256 | *cutl-28* | collagen | activated |
| Chondroitin proteoglycan synthesis |  |  |  |
| WBGene00219208 | B0365.9 | predicted chondroitin proteoglycan | activated |
| **Cuticle synthesis and molting** |  |  |  |
| WBGene00011481 | *imp-2* | intermembrane protease | activated |
| WBGene00003915 | *pan-1* | multiple functions | activated |
| WBGene00000718 | *col-145* | collagen | repressed |
| WBGene00000685 | *col-111* | collagen | repressed |
| WBGene00001074 | *dpy-13* | collagen | repressed |
| WBGene00000681 | *col-107* | collagen | repressed |
| WBGene00005017 | *sqt-2* | collagen | repressed |
| WBGene00009983 | *cut-2* | collagen | repressed |
| WBGene00001719 | *grl-10* | hedgehog ligand | repressed |
| WBGene00001694 | *grd-5* | hedgehog ligand | repressed |
| WBGene00006949 | *wrt-3* | hedgehog ligand | repressed |
| WBGene00001691 | *grd-2* | hedgehog ligand | repressed |
| WBGene00004228 | *ptr-14* | hedgehog receptor | repressed |
| WBGene00003547 | *nas-29* | metalloendopeptidase | repressed |
| WBGene00006591 | *toh-1* | metalloendopeptidase | repressed |
| **Genes associated with cuticle synthesis and molting by network associations (Kiontke et al. 2024)** |  |  |  |
| WBGene00011235 | *suro-1* | predicted metallocarboxypeptidase | repressed |
| WBGene00015841 | *skpo-2* | predicted peroxidase | repressed |
| WBGene00004120 | *pqn-32* | unknown | repressed |

An important aspect to consider about DMD-3 regulation is whether DMD-3 acts as an activator or repressor. Of the direct targets of DMD-3 that are DE in the tail tip, 97 have higher expression in WT male tails than in *dmd-3(-)* male tails and are thus activated by DMD-3 in TTM, whereas 131 have lower expression and are thus repressed by DMD-3 in TTM (Table S1). We conclude that DMD-3 can act as either activator or repressor.

### Validation of DMD-3 targets

To validate our approach to identify direct targets of DMD-3, we tested a selection of candidates from different functional categories relevant to morphogenesis: one gene encoding a transcription factor *(fos-1)*, one encoding a component of the cytoskeleton *(nmy-2),* one encoding a component of cell-cell junctions *(hmr-1)*, and one that encodes an LRR repeat-containing protein which has been implicated in a variety of functional roles (*pan-1*). The genes *fos-1, hmr-1* and *pan-1* were found to be activated by DMD-3 in the DE RNA-seq study (Kiontke, Herrera, et al. 2024); *nmy-2* was found to function in TTM via the RNAi screen (Nelson et al. 2011). Using reporter strains with endogenous fluorescent tags for these genes, we tested if deletion of the entire DMD-3 binding “peak” region and in one case only the DMD-3 associated motifs would cause at least one of two effects: a change in reporter expression in the male tail tip, or a defect in TTM.

#### fos-1

We first tested *fos-1* as an example of a transcription factor gene. This gene is involved in other morphogenetic events in *C. elegans*, e.g. in hermaphrodite anchor cell invasion and development of the uterus (Sherwood et al. 2005; Medwig-Kinney et al. 2020). We hypothesize that *fos-1* is also important for TTM because of its known role in morphogenesis and because a transcriptional reporter for *fos-1* was found to be expressed in male tail tips during TTM in a *dmd-3-*dependent manner (Kiontke, Herrera, et al. 2024). We investigated the expression of FOS-1 protein in a strain where the endogenous FOS-1a isoform is tagged with GFP ((Sherwood et al. 2005; Oommen and Newman 2007; Medwig-Kinney et al. 2020), DF400, Table 1). This reporter is expressed in the developing hermaphrodite vulva and uterus as described (Medwig-Kinney et al. 2020). In L4 males, FOS-1::GFP is expressed in the nuclei of many hypodermal cells, including in the tail tip and other cells of the male tail. It is also seen in cells of the somatic gonad. The GFP moiety does not interfere with normal FOS-1 function in the male tail tip since we observed no tail tip defects in adult males (Fig.5, n=167).

**Figure 5.**
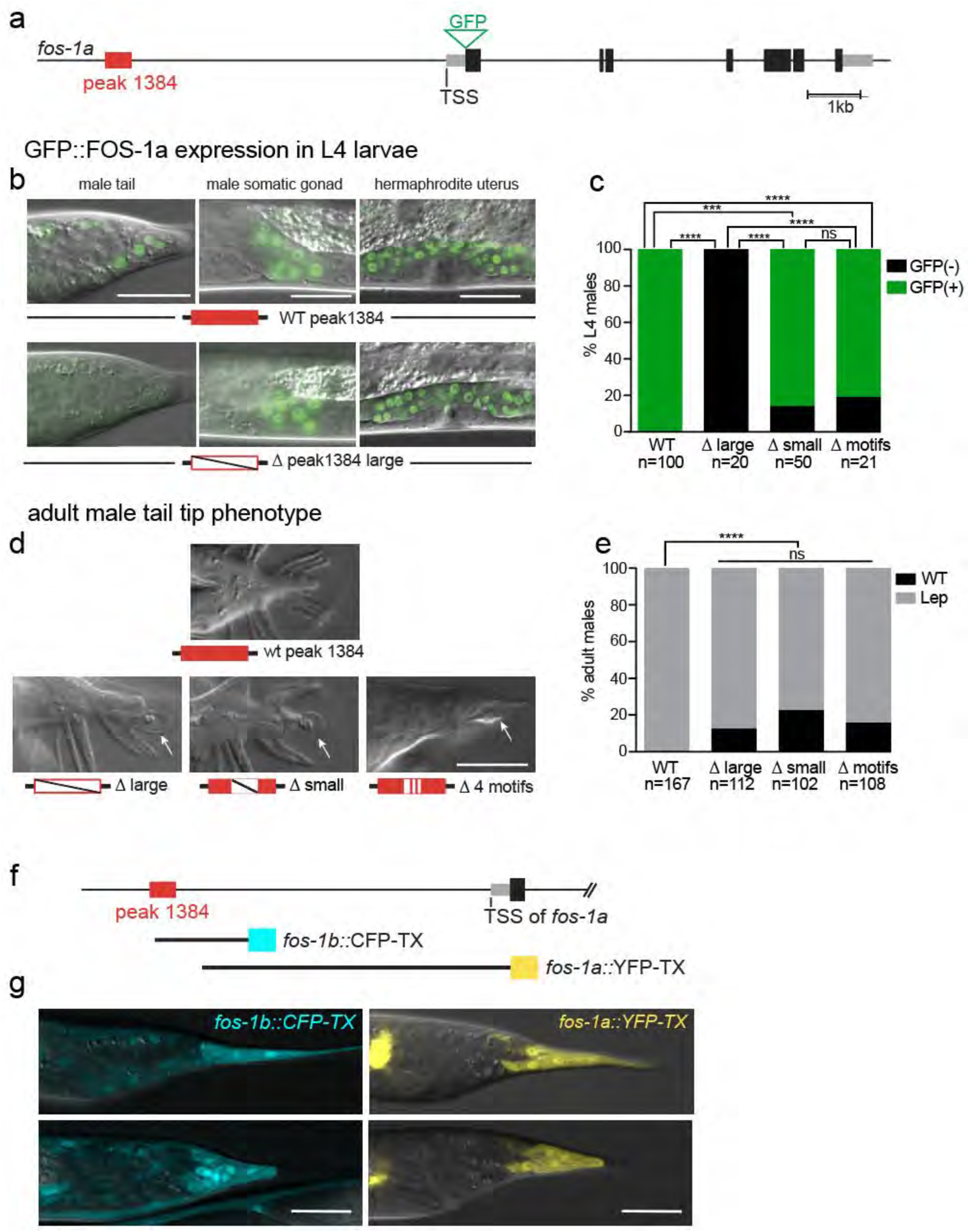
*fos-1* peak 1384 deletion mutants lose FOS-1::GFP expression in the epidermis and exhibit TTM defects. a) Gene model for the *fos-1a* locus showing the ∼400nt DMD-3-binding peak region (red box) upstream of the transcription start site (TSS) and the GFP insertion site (gray boxes = UTRs, black boxes = coding exons.) b) In L4, GFP::FOS-1 is expressed in epidermal cells including in the male tail tip and somatic gonad and in the hermaphrodite developing uterus. In peak deletion mutants, FOS-1::GFP expression is absent in epidermal cells. Expression in the somatic gonad and the hermaphrodite uterus is not affected (bottom), scale bar 20 µm. c) Quantification of male tails expressing GFP::FOS-1. d) Adult male tail tip phenotype: Some males in peak deletion and motif deletion mutants show Lep tails (arrows), indicating TTM defects. e) Quantification of males with Lep tails. f) Schematic of three transcriptional *fos-1* reporters (Sherwood et al. 2005; Kiontke, Herrera, et al. 2024). g) Expression of these reporters in male tails at two different time points during TTM. *fos-1b::*CFP-TX includes most of the peak region and is expressed in many epidermal cells and more brightly in tail tip cells. *fos-1a::*YFP-TX does not include the peak region and is only expressed in the tail tip cells.

The *fos-1* locus has a DMD-3 ChIP-seq peak (peak 1384, Fig. S2), spanning 416bp, that is 2.8kb upstream of the *fos-1b* TSS and 7.4kb upstream of the *fos-1a* TSS. Within this peak region are four instances of the DMD-3 associated motif. Deleting the entire peak region (“Δ peak1384 large”, a 477bp region containing the 416bp peak) abolished the expression of the FOS-1a::GFP reporter in all hypodermal cells of hermaphrodites and males, including in the male tail tip (100%, Fig.5, n= 20), but not in the somatic gonad of males or in the developing vulva and uterine cells of hermaphrodites. Low-penetrance (12%, n = 112) Lep TTM phenotypes also resulted (Fig. 5). When the 155-bp center of the peak region was deleted (“Δ peak1384 small”), FOS-1::GFP expression in the tail tip was lost in only 14% of males (n = 50, Fig. 5). But Lep phenotypes persisted at 22% (n = 102, Fig. 5). Finally, when just the four DMD-3 associated motif sites were excised (for a total 55-bp deletion, “Δpeak1384 motifs”), GFP expression in the tail tip was lost in 19% of L4 males (Fig. 5). Also, the penetrance of the Lep phenotype in this motif-specific mutant (15%) was similar to that for both peak region deletion mutants, indicating that the DMD-3 associated motif is sufficient to explain their effects on TTM (Fig. 5).

In summary, when the DMD-3 binding peak was deleted, we observed a significant reduction in FOS-1::GFP expression in the epidermis. The effect on expression was most pronounced in the mutant with the largest deleted region, suggesting that this entire region contains regulatory elements for *fos-1* expression in the epidermis. Deleting only the DMD-3 associated motif sites resulted in the complete absence of FOS-1 expression in 19% of the examined males. However, all deletion mutants exhibited a similar low-penetrance Lep phenotype, including those with only motif deletions, demonstrating that this region is important for TTM.

In addition to the endogenous FOS-1 protein expression, we investigated the expression of two transcriptional reporters engineered by Sherwood et al. (Sherwood et al. 2005) in L4 males. One of them (*fos-1b*::CFP-TX, teal colored in Fig. 5f and g) that likely includes the DMD-3 associated sites, shows global epidermal expression like the endogenous FOS-1a. In addition, this reporter shows clear tail tip-specific expression in L4 males. Another transcriptional reporters for *fos-1* that does not include the DMD-3 ChIP-seq peak region (*fos-1a* ::YFP-TX (Sherwood et al. 2005) show bright expression in male tail tips, but no expression in other epidermal cells (yellow in Fig. 5 f and g). This expression is dependent on *dmd-3* (Kiontke, Herrera, et al. 2024). Thus, *fos-1* expression in tail tip cells must be regulated by at least one other site in the promoter that likely binds a TF downstream of DMD-3.

#### nmy-2

Next, we tested non-muscle myosin II, NMY-2, as an example of a component of the actomyosin cytoskeleton. NMY-2 plays a role in coordinating cell shape changes and polarity during early embryogenesis in *C. elegans* and other species (Bernadskaya and Christiaen 2016; Padmanabhan et al. 2017; Seirin-Lee et al. 2020). A role for NMY-2 in TTM was demonstrated previously: RNAi against *nmy-2* results in TTM defects (Lep phenotype) and an NMY-2::mCherry transgene exhibits expression in a “cap” at the posterior end of the male tail tip during TTM (Nelson et al. 2011). We examined endogenously tagged NMY-2::mKate in L4 stage males and observed that NMY-2::mKate localized as described in the cytoplasm of the male tail and concentrated in a cap at the posterior of the retracting tail tip during TTM (Fig. 6).

**Figure 6.**
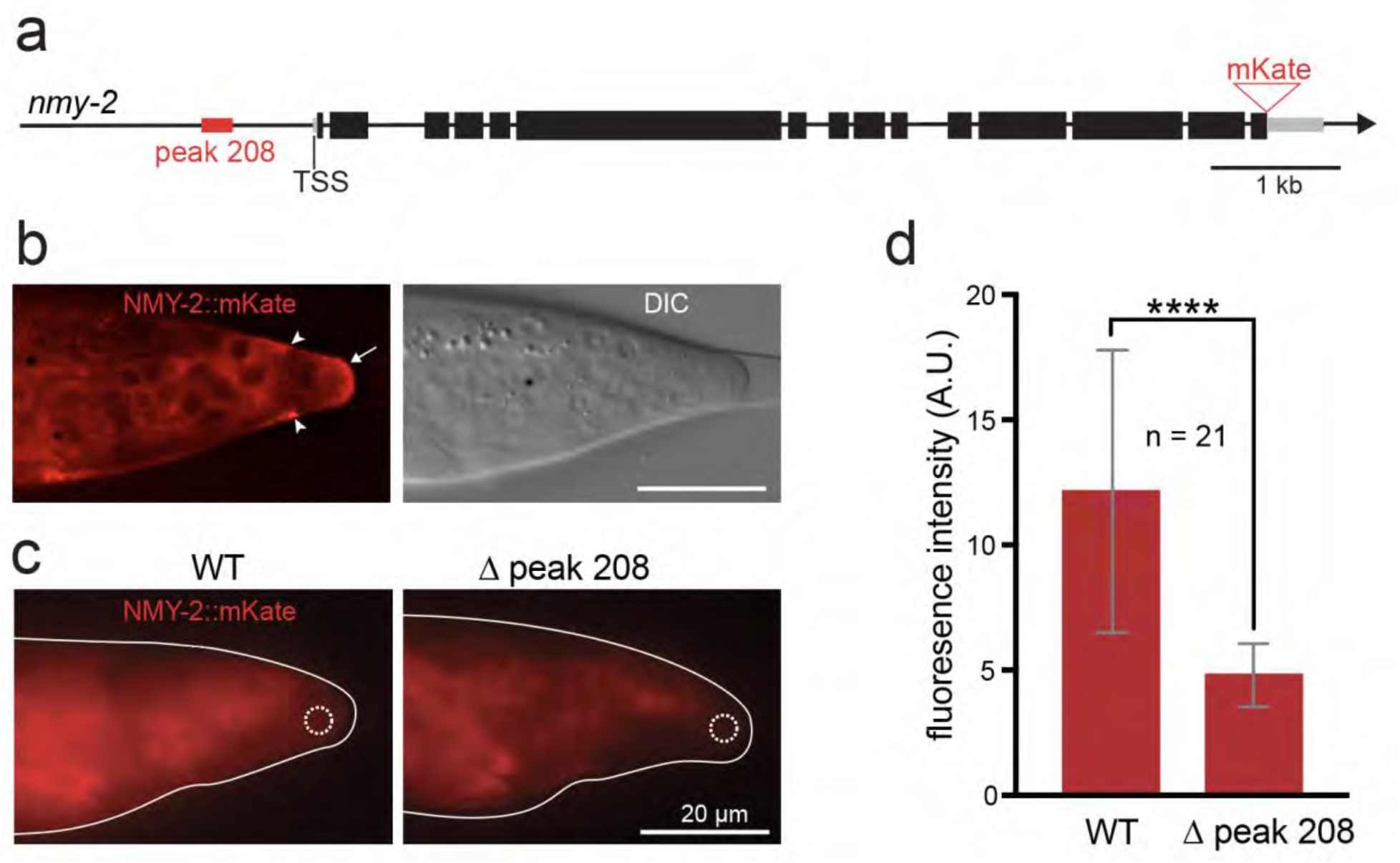
Effect of DMD-3 ChIP-seq peak deletion on the expression of NMY-2. a) Gene model for *nmy-2*; red box = 386-nt DMD-3 enrichment peak 208, gray boxes = UTRs, black boxes coding exons, red triangle = insertion site of mKate. b) Expression of NMY-2 endogenously tagged with mKate in the tail of a WT L4.4 male (left: red channel image after deconvolution and contrast enhancement, right: DIC channel image). The protein is seen in the cytoplasm of the tail. It is enriched in a cap at the end of the rounded-up tail tip (arrow) and at the adherens junctions to the tail epidermis (arrow heads). c) Non-processed red channel images of a WT male (left) and a peak deletion mutant male (right) at the L4.5 stage. The dashed white circle indicates the region of interest (ROI) used for fluorescence intensity quantification in d). d) Quantification of NMY-2::mKate mean fluorescence intensity in the ROIs of males at comparable developmental age during L4. NMY-2::mKate signal is reduced in peak mutant males compared to wildtype males; *p* < 0.0001 (t-test).

The *nmy-2* locus has a 297-bp DMD-3 enrichment peak (peak 208, Fig. S2) 737bp upstream of its TSS (Fig. 6). The FIMO motif scan identified two putative DMD-3 associated motifs, one within the assigned peak region and another 26bp away from the peak and towards the 5’ end of the coding sequence. To test the functionality of the DMD-3 binding peak, we deleted a 386-bp region encompassing the entire peak and both motif sites and looked for differences in the expression of the NMY-2::mKate protein. We found a slight but significant reduction in the intensity of the fluorescent signal (Fig. 6, *p* <0.0001) but no difference in its localization. We also investigated adult males for phenotypes indicating impaired TTM but found no defects.

#### pan-1

Next, we tested the DMD-3 enrichment peak associated with *pan-1. pan-1* was first identified in an RNAi screen for molting defects (Frand et al. 2005). The gene encodes a leucine-rich repeat protein with likely multiple functions. It was shown to be required for the proper localization of the cell-membrane-bound transcription factors MYRF-1 and MYRF-2 and their function in the synaptic rewiring of neurons (Xia et al. 2021). The protein is also localized to the membrane of intestinal cells (Gao et al. 2012) and seam cells (Xia et al. 2021), and it is associated with P granules in the germline, although it lacks an RNA-binding domain (Gao et al. 2012). *pan-1* mutants arrest their development early, but RNAi-knockdown after embryogenesis leads to arrest at the L4-to-adult transition, accompanied by defects in vulva morphogenesis and seam development (lack of alae) (Gissendanner and Kelley 2013; Xia et al. 2021). Molting and vulva development defects and expression in seam cells suggest that PAN-1 acts in the epidermis, although its function here is unclear. In the male tail tip during L4, *pan-1* mRNA expression is reduced in *dmd-3(-)* mutants relative to wild-type (File S2), indicating that the protein is transcriptionally regulated by DMD-3.

We inserted GFP at the C-terminus of PAN-1 to create a knock-in allele predicted to label all isoforms of PAN-1. We observed expression of this reporter in the tail of L3- and L4-stage males and hermaphrodites. In L3 and early L4, PAN-1::GFP is seen in large puncta at the apical surface of seam cells and at the basolateral membrane basal to the adherens junctions in hermaphrodites and males. This localization pattern is in agreement with published data for a similar allele, *syb1217* (Xia et al. 2021). In late L3 males, before the onset of TTM, we observed PAN-1::GFP at the basolateral membrane of epidermal cells in the tail. The protein is also seen in small clusters at the lateral apical surface (Fig. 7b). PAN-1 protein expression in the tail tip itself is weak, but a multicopy transcriptional reporter array with an nuclear localization signal is seen in the tail tip nuclei in males at the beginning of TTM (Fig. 7e). Early during L4 (L4.1), PAN-1::GFP transitions from the basolateral membrane of tail cells to large clusters at the apical surface of the tail, but not the tail tip. Accumulation of PAN-1::GFP is also observed in the nine Rn.aap cells on each side that undergo programmed cell death during ray development (Fig. 7c). Protein expression becomes weak later during L4. In hermaphrodite tails, PAN-1::GFP is found at the membrane of seam cells but not the tail tip cells. It is also observed in apical clusters, including in the tail region (Fig. 7d).

**Figure 7.**
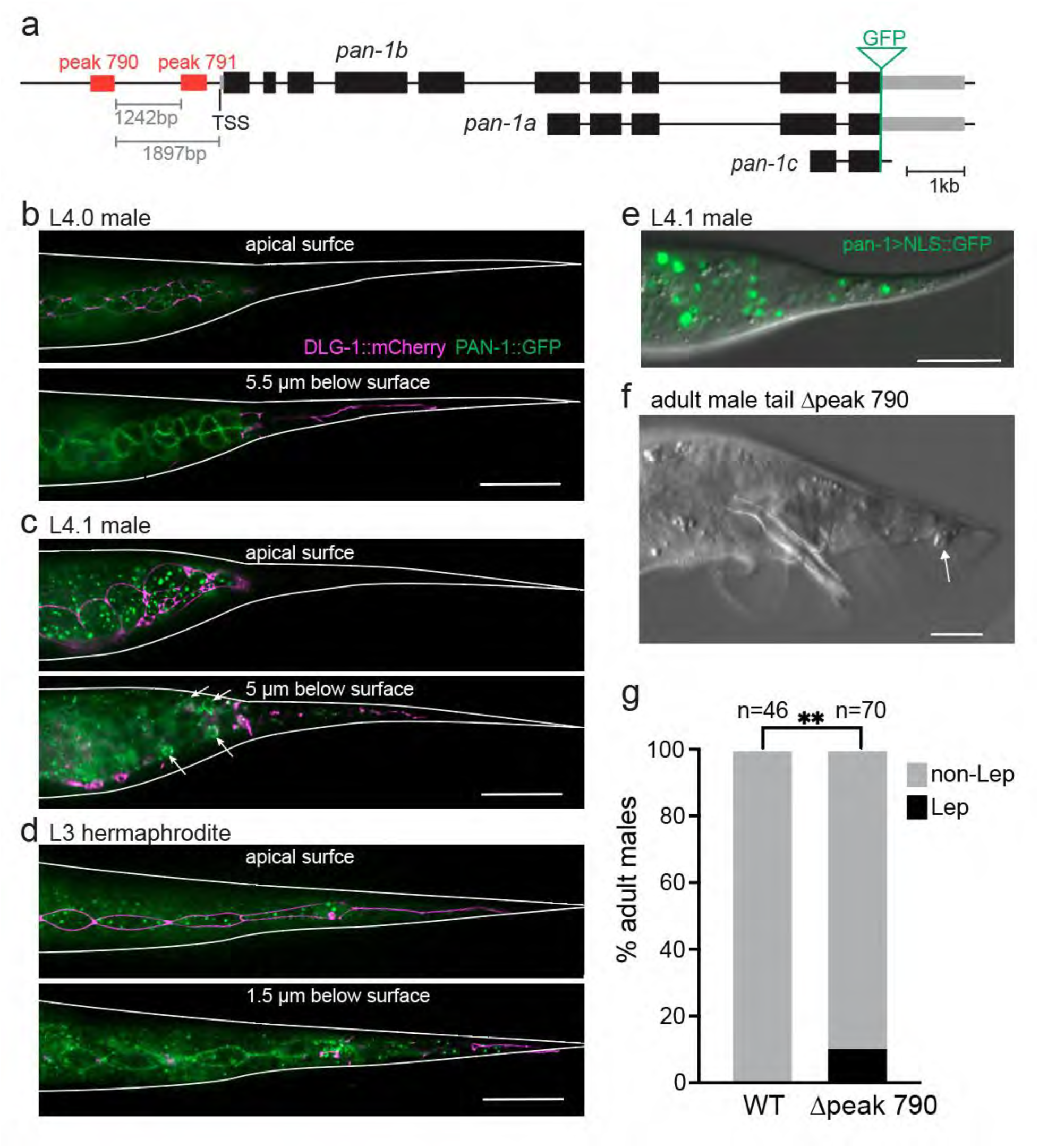
Validation of DMD-3 binding peaks in the *pan-1* enhancer region. a) Gene model for *pan-1*. Two DMD-3 ChIP-seq peaks (red boxes) are located upstream of the *pan-1b* TSS; gray boxes = UTR, black boxes coding exons, green triangle = insertion site of GFP. b) Shortly before the L3-L4 molt, PAN-1::GFP is expressed in small clusters at the apical surface in a narrow region near the seam (top) and at the basolateral membrane of ray precursor cells (bottom). No strong expression is observed in the tail tip cells. DLG-1::mCherry marks apically located adherens junctions and is used to show that the PAN-1 clusters are located apical and the PAN-1 membrane expression is basal in the epidermis. c) Expression in an L4 male at the beginning of TTM. PAN-1::GFP is now observed in large clusters at the entire apical surface of the tail except in the tail tip (top) and diffuse in the cytoplasm (bottom). The protein is also concentrated in the Rn.aap cells of the ray lineage that undergo programmed cell death (arrows). d) In L3 hermaphrodites, PAN-1::GFP is observed in clusters at the lateral apical surface (top) and at the basolateral membrane of seam cells (bottom). Some clusters are also seen in the tail tip. e) A transcriptional reporter for *pan-1* with a nuclear localization signal is expressed in the tail tip. f) Adult male tail tip in a peak 790 deletion mutant with small Lep tail tip (arrow). g) Quantification of the Lep phenotype in peak 790 deletion mutants; p = 0.0269 (chi-square test).

Two DMD-3 ChIP-seq peaks are situated 5’ to the TSS for the longest isoform, *pan-1b* (Fig. S2). The first peak (790) spans 325bp, contains seven DMD-3 associated motif sites and is located 1.9kb upstream of the TSS. The second peak (791) spans 311bp, contains also seven DMD-3 associated motif sites and is located 344bp upstream of the TSS. These peaks are separated by 1.2kb (Fig. 7). We first deleted 340bp containing the entire peak 791. Careful inspection of PAN-1::GFP expression during TTM and the phenotype of adult male tails revealed no effect of this mutation on either PAN-1::GFP expression or TTM phenotype. Next, we deleted peak 790. Again, we did not see any difference in PAN-1::GFP expression. However, with this mutation, we observed Lep tails in 10% of adult males, indicating defective TTM (Fig. 7f, n = 70, *p* = 0.0269).

Deleting both peaks resulted in severe molting defects, worms with a ’clear’ appearance, and larval arrest (Fig. S3). These phenotypes were previously described for the *pan-1(gk142)* mutant allele that is assumed to be a molecular null, as well as for *pan-1* RNAi treatment by injection or soaking (Gao et al. 2012; Gissendanner and Kelley 2013; Xia et al. 2021). Our results thus suggest that both peak regions are involved in *pan-1* regulation, but only peak 790 plays a specific role in TTM.

**Figure S3.** Deleting both peaks at the pan-1 locus causes global phenotypes resembling the phenotype of a pan-1 loss-of-function mutant. Worms displayed larval arrest at different stages, were not able to molt and had clear appearance (A). (B) A tiny worm with an L4 gonad. (C) Many worms remained tiny.

#### hmr-1

HMR-1 is a *C. elegans* classical cadherin. It participates in the catenin/cadherin complex at epithelial junctions, is required for cell-cell adhesion and is involved in several morphogenetic processes in the embryo, e.g. gastrulation by apical constriction, ventral enclosure and elongation (Hardin et al. 2013). HMR-1 also plays a role in polarity establishment in early embryos (Klompstra et al. 2015) and in the developing intestine (Naturale et al. 2023). In one-cell embryos, HMR-1 has a junction-independent role in stabilizing the actomyosin cortex during cytokinetic furrow formation (Padmanabhan et al. 2017). *hmr-1* mRNA levels are elevated in the tail tip of WT males relative to *dmd-3* mutant males, suggesting that DMD-3 activates *hmr-1* expression (Kiontke, Herrera, et al. 2024). To investigate HMR-1 expression during TTM, we took advantage of an endogenous C-terminal GFP fusion to HMR-1 that marks all isoforms of the protein (Marston et al. 2016), and investigated its expression in male tails during TTM. As expected, HMR-1 is seen at apical junctions of seam cells and developing ray precursor cells. However, in early L4 males (L4.1), HMR-1 has disappeared from junctions of the tail tip cells, although other components of adherens junctions (AJM-1 and DLG-1) are still present (Fig. 8b), and the junctions are visible in EM sections at this time and only fully disassemble later during TTM (Nguyen et al. 1999; Kiontke, Fernandez, et al. 2024). At that time (L4.3) HMR-1 foci appear at the surface of the tail tip cells and the body epidermis hyp7 (Figure 8b middle panel). When *dmd-3* is mutated, HMR-1 remains localized to the junctions between tail tip cells, which do not disassemble in *dmd-3(-)* males. The *hmr-1* gene encodes 5 isoforms, the largest, *hmr-1b* leading to a 9485nt long transcript with 31 exons. This protein was shown to specifically affect axonal patterning (Broadbent and Pettitt 2002).

**Figure 8.**
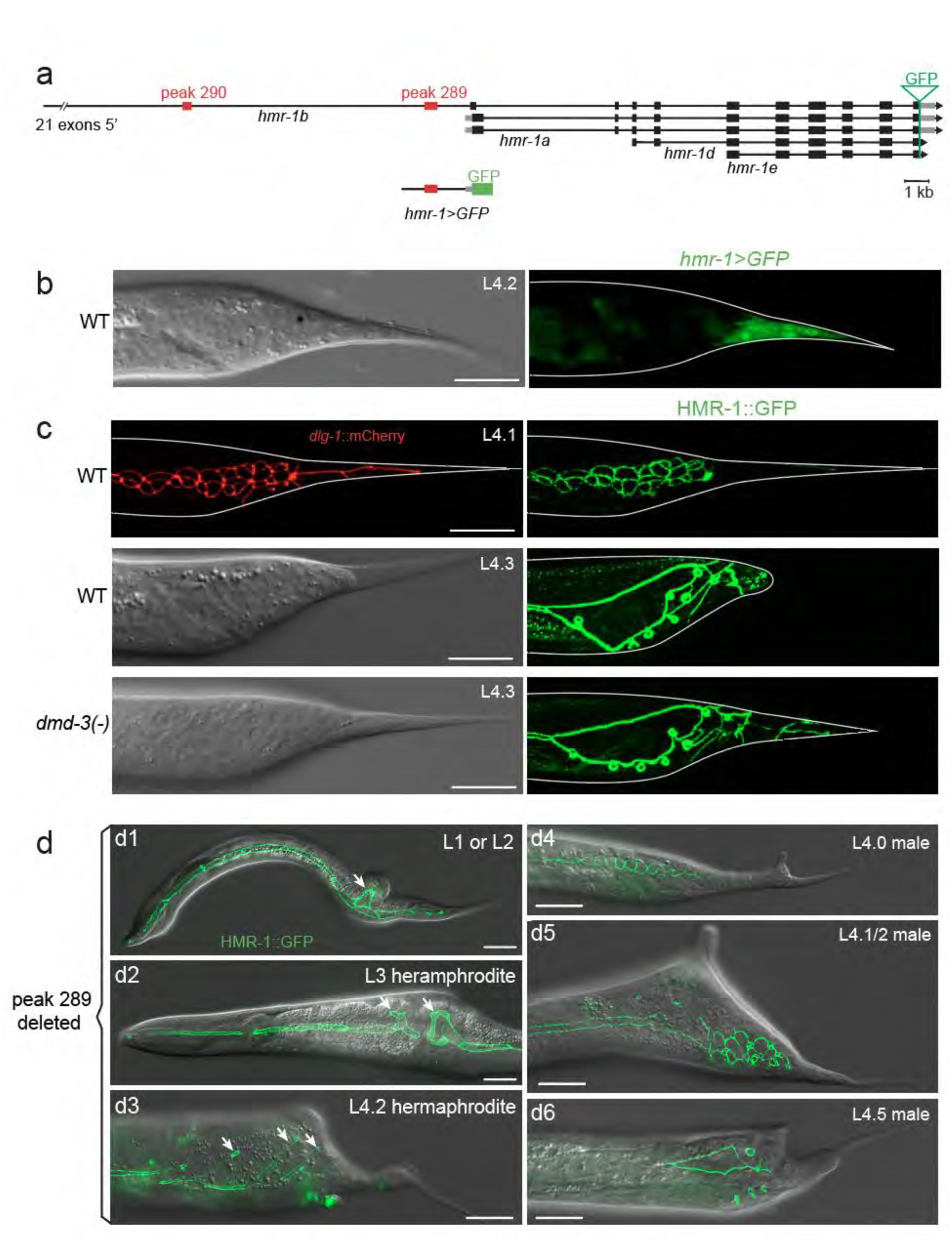
Expression of *hmr-1* in the male tail and effect of deleting a DMD-3 ChiP-seq peak. a) Gene model for *hmr-1*, the 5’ part of isoform b is truncated in the intron upstream of exon 22. red boxes = DMD-3 enrichment peaks 289 and 290, gray boxes = UTRs, black boxes coding exons, green triangle = insertion site of GFP; this fusion labels all isoforms. Below is a schematic of the transcriptional reporter with a region 2620nt upstream of the TSS of *hmr-1a* driving GFP. b) An L4.2 male expressing the transcriptional reporter for *hmr-1a* in the tail tip cells (right), left panel shows DIC channel. c) expression of the endogenous HMR-1::GFP fusion protein in L4 WT males and one *dmd-3(-)* male. Top row: in L4.1 males, adherens junctions are not yet disassembled as indicated by DLG-1::mCherry expression (left), but HMR-1::GFP is not seen at the junctions (right). Middle row right: in L4.3 males whose tail tip has rounded up (DIC channel left), HMR-1::GFP is expressed in puncta at the surface of the tail tip cells. Puncta are also observed in the body epidermis; left: DIC channel. Bottom row right: L4.3 males mutant for *dmd-3* continue to show HMR-1::GFP in the tail tip cells that have not rounded up and have not disassembled their adherens junctions left: DIC channel. d) Examples of the phenotype seen after deleting peak 289. d1) Young larva showing branched seam cells (arrow) and dorsal bulge in the posterior body region. d2) Anterior body region of an L3 hermaphrodite with branched seam cells and dorsal bulge (arrow). d3) L4 hermaphrodite with a dorsal epidermal bulge in the tail region and multiple branched seam cells (arrows). d4-6) Three males at different stages during L4 showing dorsal epidermal bulges in the tail region. All scale bars = 20µm.

Of the other isoforms, *hmr-1a* encodes the protein present in the junctional cadherin/catenin-complex. A transcriptional reporter for this isoform (Fig. 8a) shows enrichment of expression in the tail tip of males at the beginning of TTM (Fig. 8c), suggesting that this isoform is functional in TTM. Our observations indicate that in TTM, HMR-1 acts in a junction-independent role: the transcriptional reporter and RNA-seq data show that *hmr-1* transcription is activated by DMD-3 at the beginning of TTM, even though the protein disappears from the adherens junctions in the tail prior to their disassembly. It later shows up in puncta. The function of these HMR-1 foci in TTM is unknown.

Two ChIP-seq peaks are located in an intron of *hmr-1b* upstream of the all other isoforms: peak 289 lies 1168nt upstream of the TSS of *hmr-1a* and is 554nt long, peak 290 lies 11105nt upstream of this TSS and encompasses 350nt. We deleted peak 290 in the HMR-1::GFP strain and did not see any effect on protein expression or adult male tail tip phenotype. When we deleted peak 289, we observed global morphology defects throughout development. Many males and hermaphrodites at all stages display dorsal epidermal bulges in various body regions (Fig. 8d), which are accompanied by seam cell division and polarity defects. Seam cells can be severely disrupted, showing gaps and branchings (Fig. 8d right panel). This phenotype is seen in 58% of hatched L1 (n=83). The defect causes some larval lethality which explains why 30% of adults (n=101) were affected. Dorsal epidermal bulges have been previously described for the small fraction of *hmr-1-*null mutant animals that do not arrest in embryogenesis due to ventral enclosure defects (Costa et al. 1998). Dorsal epidermal bulges are more frequently observed in animals mutant for the catenins *hmp-1* and *hmp-2* which, however, also die as embryos or larvae. Deleting the entire peak 289 led to a comparatively mild defect in the junctional catenin/cadherin complex, which indicates that the region of peak 289 contains regulatory elements for *hmr-1* expression in the entire epidermis. Because of the relatively high penetrance of general epidermal defects in the peak deletion mutants, we were unable to evaluate adult males for TTM defects, which we expected to observe in only a small fraction of males. In L4 males, we did not see any obvious difference in protein expression, but this could also be rare and subtle. How DMD-3 binding at this enhancer affects TTM is therefore currently unknown.

## DISCUSSION

### Summary of results

ChIP-seq in L4 males found 1755 enrichment peaks corresponding to 6061 genes (in a 10kb window around the TSS), any of which are potentially regulated by DMD-3. 270 of these genes were known to function in the male tail during TTM from previous TTM-specific studies (Nelson et al. 2011; Kiontke, Herrera, et al. 2024). We identified a DMD-3-associated binding motif, which is strikingly similar to the motif found for EOR-1. A transgenic array tiled with this motif but not a control array caused TTM defects, presumably because it sequestered DMD-3 away from TTM genes. We validated DMD-3-bound regions as playing a role in TTM regulation for three genes: For *fos-1* and *pan-1,* deletion of this region leads to TTM defects. For *nmy-2,* deletion of the region results in diminished expression of an endogenous reporter.

### How does DMD-3 work as a master regulator of TTM?

We generated a dataset of direct and indirect targets of DMD-3 for TTM, which allows us to evaluate the hypotheses for the role of DMD-3 as a master regulator of this morphogenetic process. One hypothesis is that the master regulator sits at the top of a hierarchy of TFs and that most or even all of its targets are other TFs that control the machinery of morphogenesis. Alternatively, the master regulator TF may target genes of this machinery directly.

We found 270 direct targets of DMD-3, and even though these are enriched in TFs (28), they contain many other types of genes, including components of the cellular machinery executing TTM. Thus, the GRN of TTM is indeed hierarchical with DMD-3 regulating TFs that must target the other TFs which are not direct DMD-3 targets. DMD-3 also directly targets genes for “effectors” such as intracellular components of signaling pathways and protein modification enzymes (Kiontke, Herrera, et al. 2024) and genes for proteins in modules responsible for the execution of TTM. Notably, among the direct targets of this TF are genes that encode components of universal modules, e.g. components of the cytoskeleton, junctions, vesicular transport, metabolism and several proteins acting in the unfolded protein response. Several of these direct targets of DMD-3 affect TTM when knocked down by RNAi but are not differentially expressed in *dmd-3(-)* versus wild-type genotypes. This may be expected for universally acting genes, which may be redundantly regulated by several TFs; the role of DMD-3 in this case may be to ensure their robust expression in tail tip cells. Several “conditional modules”— groups of genes that act together in TTM but are not universally active in every cell— had been identified previously (Kiontke, Herrera, et al. 2024). Two of these modules, chondroitin proteoglycan synthesis and the hyperosmotic stress response, which are activated during TTM, are not direct DMD-3 targets. Likewise, cuticle synthesis or maintenance, a module comprising at least 43 genes repressed upon TTM onset, is mainly controlled indirectly. We conclude that the TTM GRN downstream of DMD-3 is tiered with a hierarchical cascade of TFs (Fig. 9) and modular.

**Figure 9.**
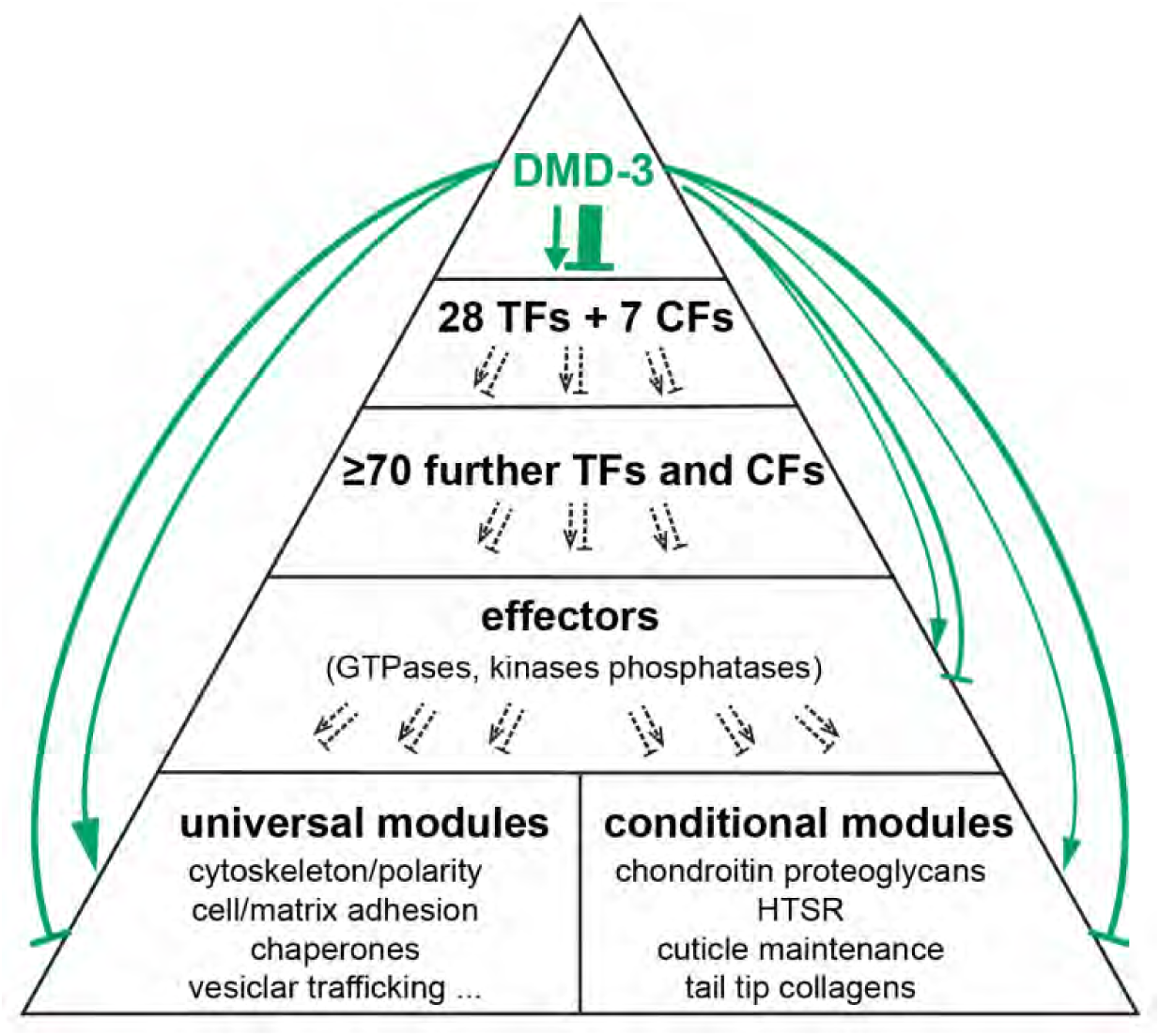
Gene-regulatory network model for TTM downstream of DMD-3. DMD-3 directly regulates several TFs and their CFs, but previous work has shown that at least twice as many other TFs and CFs are downstream of DMD-3 and are thus indirect targets of DMD-3. We posit that TFs mostly regulate effectors which then regulate the genes that execute TTM. The latter are organized in several modules of genes acting together. Universal modules are groups of proteins that are universally active in the cells, conditional modules proteins of genes that act together but are not always expressed.

### How does DMD-3 work as a TF?

Our data, together with published information, give some indication as to how DMD-3 functions as a TF at the molecular level. Three pieces of information suggest that DMD-3 may not bind DNA directly but acts together with another TF. (1) The enrichment values for DMD-3 in our ChIP-seq experiments were relatively low compared with experiments with other TFs (e.g. Berkseth et al. 2013). Although this could be due to low DMD-3 protein abundance in our samples, it might also indicate that DMD-3 binds DNA with low affinity. (2) A previous study that analyzed DNA binding motifs of *C. elegans* TFs with protein-binding microarrays (Narasimhan et al. 2015) could not find evidence for a strong binding motif for DMD-3. The weak motif predicted by this study (consensus AATGTTTC) is similar to one detected using yeast-one-hybrid analysis (consensus WWTGTTTCAAWW) (Fuxman Bass et al. 2014), but it does not match a binding motif previously predicted for DMD-3 bioinformatically (TGTAACA) (Mason et al. 2008; Siehr et al. 2011) or the DMD-3 associated motif identified and validated by us (TTCTCTGCGTCTCTC). It is possible that this motif is not actually bound by DMD-3 but instead by another TF with which it cooperates. (3) Deletion of the DMD-3 associated motifs in the cis-regulatory region of *fos-1* abolished the expression of *fos-1* in the epidermis, but no such effect was seen in *dmd-3(-)* mutants. This suggests that DMD-3 is either not the only TF involved in binding this motif, or DMD-3 does not bind to DNA at all but instead interacts with another TF that does.

One of the factors with which DMD-3 cooperates might be EOR-1. This is because the DMD-3 associated motif has high similarity to the binding motif of EOR-1 (Kumar et al. 2015; Daugherty et al. 2017) (Fig. 3), and 42% of the DMD-3 ChIP-seq enrichment peaks overlap with an enrichment peak for EOR-1. Some of the remaining DMD-3 binding sites might also bind EOR-1, given that the experimental conditions for the ChIP-seq experiments were different. Specifically, we used L4 males, whereas the EOR-1 data was obtained for hermaphrodite and mixed-sex samples at the L3 stage. Alternatively, DMD-3 might associate with other TFs at these sites as well as at the 21% of DMD-3 ChIP-seq peak regions that do not contain the motif. EOR-1 is a TF that is hypothesized to regulate chromatin accessibility or chromatin state (Daugherty et al. 2017; Shinkai et al. 2018) and might act in a complex with other transcription factors (Kumar et al. 2015; Daugherty et al. 2017). EOR-1 plays, together with EOR-2, a role in the development of the nervous system, including sensory structures in the male tail (Hoeppner et al. 2004), and in vulva and excretory system development (Howard and Sundaram 2002). A role for EOR-1 in TTM is in line with these observations.

The differential expression data furthermore indicate that DMD-3 functions as both a repressor and an activator. These opposing functions are likely not due to differences in the binding site sequence as reported for some TFs (Latchman 2001), because there is no enrichment for one DMD-3 binding motif versus another at genes that are positively or negatively differentially expressed. Instead, DMD-3 is more likely to interact with a co-activator/co-repressor partner (Latchman 2001; Valin and Gill 2007). Combined with the evidence above, we suggest that the molecular function of DMD-3 is to act as a mediator between TFs that bind directly to DNA, co-activators and co-repressors, and the transcription machinery.

### How are DMD-3-associated sites involved in gene regulation?

Deleting the putative DMD-3 binding sites in the endogenous cis-regulatory regions of four candidate genes yielded insights into the role of these sites for the regulation of the genes. For five of the six DMD-3 associated peak sites that we deleted, a resulting tissue-specific change in protein expression level and/or a defect in TTM indicate that they contain tissue-specific enhancers.

The strongest response was observed for *fos-1*, in which deletion of peak 1384 led to a complete absence of the epidermal expression of the FOS-1 protein in males and hermaphrodites. Thus, this peak contains an enhancer element for epidermal expression. Deleting only the DMD-3 associated motif sites had the same effect (albeit not with 100% penetrance). As outlined above, we suggest that EOR-1 and not DMD-3 may bind these motif sites directly. EOR-1 was hypothesized to regulate chromatin accessibility for other TFs (Daugherty et al. 2017) and may thus be associated with another TF. This TF cannot be DMD-3 in the epidermal tissues where *dmd-3* is not expressed. That is, this “epidermal enhancer” conveys global epidermal *fos-1* expression, but DMD-3 is not expressed in hermaphrodites or globally in the epidermis of males. Thus, another TF (TF1 in Fig. 10a) must associate with EOR-1 to drive *fos-1* expression in the epidermis.

**Figure 10.**
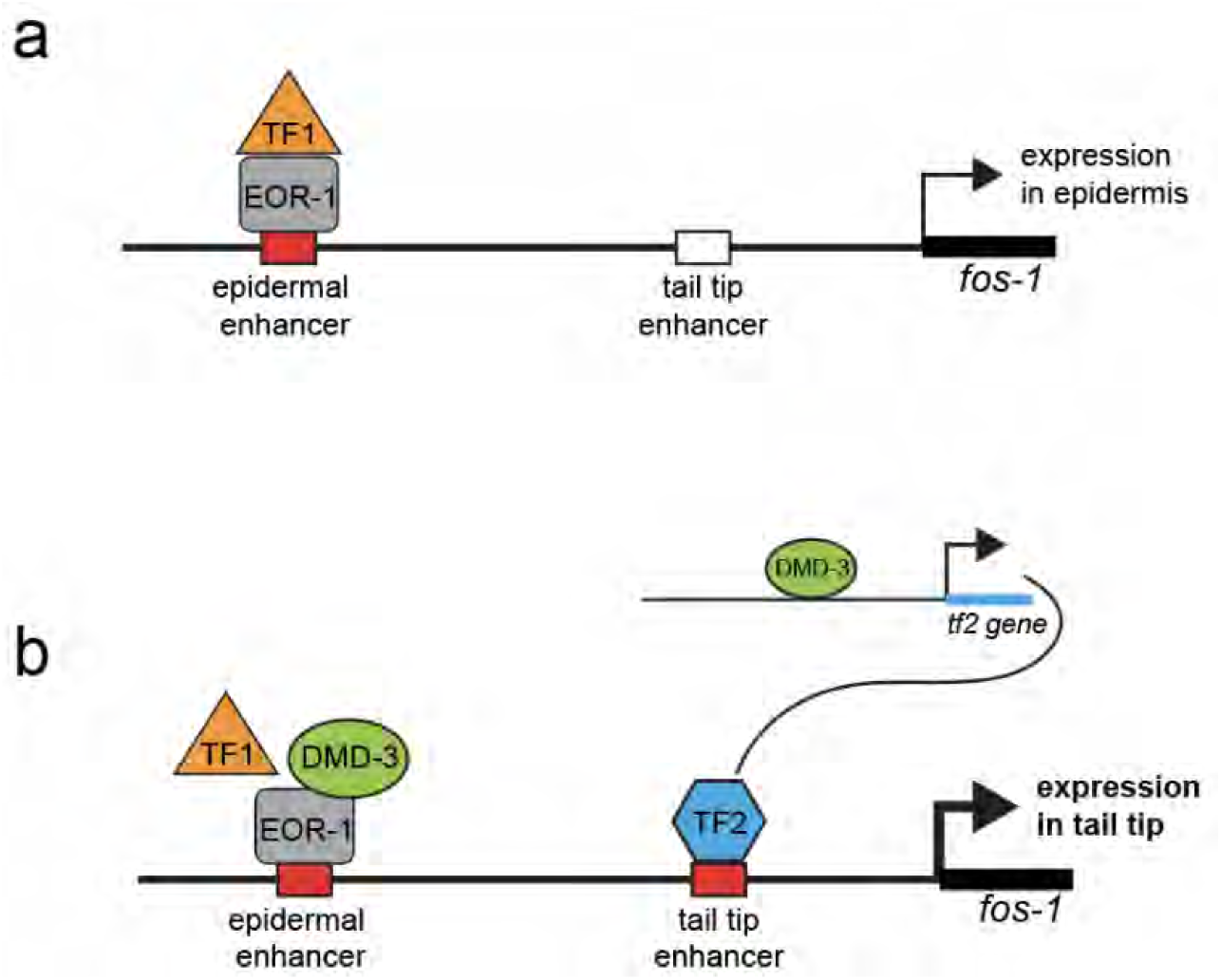
Model for the transcriptional regulation of *fos-1*. a) In all epidermal cells, including the male tail tip before TTM, the enhancer in the cis-regulatory region of *fos-1* at peak 1384 is bound by EOR-1, which may modify chromatin to make the site accessible for TF1, which drives expression of *fos-1* in all epidermal cells. The tail tip enhancer is unbound. b) In tail tip cells, at the beginning of TTM, DMD-3 binds to EOR-1 at the epidermal enhancer either together with TF1 or instead of it. DMD-3 also drives the expression of TF2, which binds to the tail tip enhancer in the cis-regulatory region of *fos-1;* an increase of *fos-1* expression in the tail tip ensues.

In certain male epidermal cells, e.g. the tail tip, DMD-3 either associates with this hypothesized complex or replaces the putative epidermal TF. This would explain why in the male tail tip, *fos-1* mRNA expression is upregulated in a *dmd-3-*dependent manner (Kiontke, Herrera, et al. 2024), and that deletion of the DMD-3 associated motifs at the epidermal enhancer results in TTM defects. Expression of one of the transcriptional reporters (*fos-1b*::CFP-TX) that we analyzed is consistent with this explanation. Another reporter, however, does not include the peak 1384 region or indeed any DMD-3 binding site (*fos-1a*::YFP-TX, gbyEx331) yet shows a clear tail-tip-specific expression that is *dmd-3* dependent. Thus, a second enhancer element must be present that mediates *dmd-3*-dependent *fos-1* upregulation only in the tail tip. Because there are no other DMD-3 binding peaks in the cis-regulatory region of *fos-1,* a TF downstream of DMD-3 must bind to this element to convey tail-tip-specific expression of *fos-1* (TF2 in Fig. 10b). Where this element is located and what TF binds there is currently unknown. This indicates that there is a feed-forward circuit in the regulation of *fos-1* by DMD-3 to ensure strong expression in the tail tip.

For the other candidate DMD-3 target genes that we tested, deleting DMD-3 ChIP-seq peaks had more subtle effects but still showed that these peaks contain enhancers. Deletion of peak 208 5’ of *nmy-2* led to a slight reduction of cytoplasmic NMY-2 protein expression and no TTM defect. TTM is affected when *nmy-2* mRNA levels are severely reduced by RNAi (Nelson et al. 2011). It is possible that a threshold level of NMY-2 is required for TTM and that DMD-3 binding to the enhancer of *nmy-2* might boost expression of the gene in the tail tip to ensure above-threshold levels. This may thus contribute to the robustness of TTM.

We could not detect any change in PAN-1 protein expression upon deletion of the two DMD-3 ChIP-seq peaks 5’ of its gene individually. However, we found that deleting both peaks together phenocopied the *pan-1* loss-of-function mutation and thus confirmed that these regions contain enhancers for *pan-1* regulation. We also observed male tail defects when we deleted peak 790, indicating that DMD-3 binding to this region plays a role in TTM. We did not see changes in protein expression, which may be due to the already low protein level in the tail tip. A small decrease may be below our detection capability. The *pan-1* mRNA level in the tail tip was also only slightly higher in wild type males than in *dmd-3* mutant males (log2-fold change 1.2).

Based on the phenotypic effects on seam cell and epidermal morphology, deletion of the DMD-3 ChIP-seq peak 289 in the cis-regulatory region of *hmr-1* uncovered an epidermal or seam-cell-specific enhancer. Since we observed L1 worms with seam cell defects, this enhancer must already be active in embryogenesis. These early defects appear to lead to a dorsal hump phenotype known from worms mutant in catenin-cadherin complex members HMP-1 and HMR-1. How binding of DMD-3 to this region may affect TTM could not be fully evaluated, but any potential effect must be subtle since we did not see any obvious tail tip defects.

### Conclusions

In conclusion, although DMD-3 is a master regulator of sexually dimorphic TTM, it likely interacts with other factors that provide DNA-binding and co-activator/repressor activities. Specifically, it may piggyback on other factors like EOR-1, which already recognizes sites in tissue-specific enhancers. Additionally, TFs regulated downstream of DMD-3 in the GRN hierarchy govern genes. In the case of fos-1, DMD-3 and a downstream TF both appear to participate in a feed-forward circuit to boost expression tail tip-specifically. Thus, future work is needed to identify this other TF and the tail tip-specific cis-regulatory element it recognizes. More broadly, understanding how DMD-3 is connected to the cellular machinery of morphogenesis will entail the delineation of the TF network and targets of these downstream TFs. There is evidence that *dmd-3* itself is a feedback target of itself, *mab-3*, and perhaps other genes (Mason et al. 2008; Nelson et al. 2011). That is, the “master regulator” of TTM may have to be more broadly defined as a network, such as that suggested for *Drosophila* eyeless (Davis and Rebay 2017). Related to the architecture of networks regulating morphogenesis, it has also been proposed that modularity is an important, though not exclusive organizing principle (Davies, 2023). Thus, an important question to address in future work is the degree to which the cellular processes are governed by separate or overlapping TFs.

## Data availability

Raw and processed data files are deposited at GEO and are available through GEO accession number GSE282566. Scripts for the ChIP-seq analysis can be found at (https://github.com/classymagpie/DMD3_ChIPseq)

Files S1 (folder) contains data from the motif enrichment analyses with HOMER and MEME. File S2 contains a list of differentially expressed genes from (Kiontke, Herrera et al. 2024) with a adjusted p-value threshold of 0.05. Files S3 (folder) contain location of the DMD-3 ChIP enrichment peaks and gene assignments.

Strains can be obtained from the authors upon request; some strains will be deposited at the *Caenorhabditis* Genetics Center.

## Acknowledgements

We thank Alyssa Woronik for assistance with bioinformatics.

Strain FT1838 was kindly provided by Jeremy Nance; other strains were provided by the CGC, which is funded by NIH Office of Research Infrastructure Programs (P40 OD010440).

## Supplemental Material

File S1. De novo motif scan output from HOMER and MEME-ChIP, easy access motif, and FIMO output.

File S2. Reanalysis of the dataset from Kiontke et al. (Kiontke, Herrera, et al. 2024) with an adjusted p-value threshold of 0.05 instead of 0.01 that was used in the original study.

File S3. Gene assignments by genomic distance of ChIP-seq enrichment peak relative to TSSs (using 2, 4, 6, 8, and 10 kb windows). Statistics and DMD-3-associated binding motif locations pertaining to the 1755 DMD-3 ChIP-seq peaks.

Table S1. Table of DMD-3 target genes that overlap with either RNA-seq (Kiontke, Herrera, et al. 2024) or TTM RNAi screen (Nelson et al. 2011).

